# Intercellular crosstalk regulating ARRB2/RARRES1 is involved in transition from fibrosis to cancer

**DOI:** 10.1101/2021.09.08.458161

**Authors:** Robert Schierwagen, Peter Dietrich, Judith Heinzen, Sabine Klein, Frank E. Uschner, Cristina Ortiz, Olaf Tyc, Sandra Torres, Christoph Hieber, Nico Kraus, Richard T. Premont, Leon D. Grünewald, Johanne Poisson, Pierre-Emmanuel Rautou, Glen Kristiansen, Jordi Gracia-Sancho, Marko Poglitsch, Isis Ludwig-Portugall, Thomas Walther, Christian Trautwein, Zeinab Abdullah, Christian Münch, Christoph Welsch, Mercedes Fernandez, Stefan Zeuzem, Richard Moreau, Claus Hellerbrand, Krista Rombouts, Wolfgang Kastenmüller, Anna Mae Diehl, Jonel Trebicka

## Abstract

Progressive fibrogenesis in chronic liver injury is often associated with cancer development. Beta-arrestin-2 (ARRB2) is a regulator of the profibrotic Angiotensin II type 1 receptor (AGTR1). The role of ARRB2 in liver fibrosis and in the transition from fibrosis to cancer is not fully understood and was investigated in this study.

This study demonstrates that upregulation of the retinoic acid receptor responder 1 (RARRES1) in HSC mediated by ARRB2 leads to fibrosis. This process is driven by exosomal ARRB2 transfer to HSC, major fibrosis contributors, from injured hepatocytes, which highly express ARRB2. By contrast, downregulation of RARRES1 in hepatocytes induces malignant transformation and hepatocellular carcinoma (HCC) development. Consequently, Arrb2-deficient mice show higher number and size of liver tumors than wild-type mice in a hepatocellular carcinoma model with fibrosis. The identified relationship between ARRB2 and RARRES1 was observed in at least two species, including human cells and tissues in fibrosis and HCC and has a predictive value for survival in cancer patients. This study describes the discovery of a novel molecular pathway mediating the transition from fibrosis to cancer offering potential diagnostics and therapeutics.

## Introduction

Chronic tissue injury leads to fibrosis in many organs (1). Liver fibrosis is a good model to study fibrogenesis in response to chronic injury (2) and represents a premalignant condition associated with liver cancer. Yet, the relationship between fibrosis and cancer is not well understood. For example, retinoic acid receptor responder 1 (RARRES1), which is a putative tumor suppressor, with decreased expression in tumor, is one out of only two markers upregulated in progressive fibrosis of liver, kidney and lung (3). To understand the relationship between fibrosis and cancer, a deeper look into fibrosis is required. The major contributors to liver fibrosis are activated transdifferentiated hepatic stellate cells (HSC) (4). In those cells, profibrotic properties have been attributed to angiotensin II receptor type 1 (AT1R) via G protein mediated downstream signaling (5, 6).

However, the AT1R signaling is regulated by beta-arrestin-2 (ARRB2), which is a soluble cytoplasmic ubiquitously expressed protein. Thereby, ARRB2 can interact with the AT1R in multiple ways. On the one hand, ARRB2 desensitizes the G protein signaling downstream of the AT1R and thereby restores homeostasis in tissue (7, 8). One way of AT1R desensitization is internalization of the receptor which is facilitated by ARRB2. On the other hand, ARRB2 also promotes distinct, G protein-independent signaling downstream of receptors such as the AT1R, through, amongst others, mediators of the mitogen-activated protein kinase (MAPK) pathway (9). Previous reports have associated beta-arrestin-2 and the closely related beta-arrestin-1 with pulmonary fibrosis (8) and atherosclerosis (10). Furthermore, it has been described that beta-arrestin-1 is involved in liver cancer development (11).

Here, we describe the role of ARRB2 in liver fibrogenesis and uncover a novel intercellular crosstalk between parenchymal cells (hepatocytes) and extracellular matrix-producing cells (HSC) in the liver. Furthermore, we propose a molecular basis involving ARRB2 and RARRES1 for the development of cancer in liver fibrosis and provide first hints for a broader importance of the discovered cascade in carcinogenesis by identifying homologues expression patterns in the kidney for renal fibrosis and cancer development.

## Results

### Inhibition and stimulation of beta-arrestin-2 in angiotensin II receptor type 1-mediated liver fibrogenesis

AT1R plays a pivotal role in fibrosis. To assess the role of ARRB2 in AT1R-mediated fibrosis, knock-out mice for AT1R (*Agtr1*) and *Arrb2* as well as double knock-out mice for *Agtr1*/*Arrb2* were used. Liver fibrosis was induced by bile duct ligation (BDL) as a model for cholestatic fibrosis, and by carbon tetrachloride (CCl_4_) intoxication as a model for toxic fibrosis. In this setting, accumulation of hepatic extracellular matrix (ECM) was quantified by hydroxyproline, a major component of collagen. While *Agtr1* deficiency decreased hepatic fibrosis in both models as previously described (5), *Arrb2* deficiency showed a slightly stronger protective effect than *Agtr1* knock-out. Mice with a double knock-out of *Agtr1* and *Arrb2* developed a similar severe fibrosis as *Arrb2* deficient littermates, suggesting ARRB2 seems to be the key mediator of AT1R-associated fibrosis (Figure 1A). Reduced collagen production in *Arrb2*-deficient mice was confirmed at the level of gene expression (Figure 1B) and at the protein level by Sirius red staining and quantification thereof (Figure 1C).

**Fig. 1.**
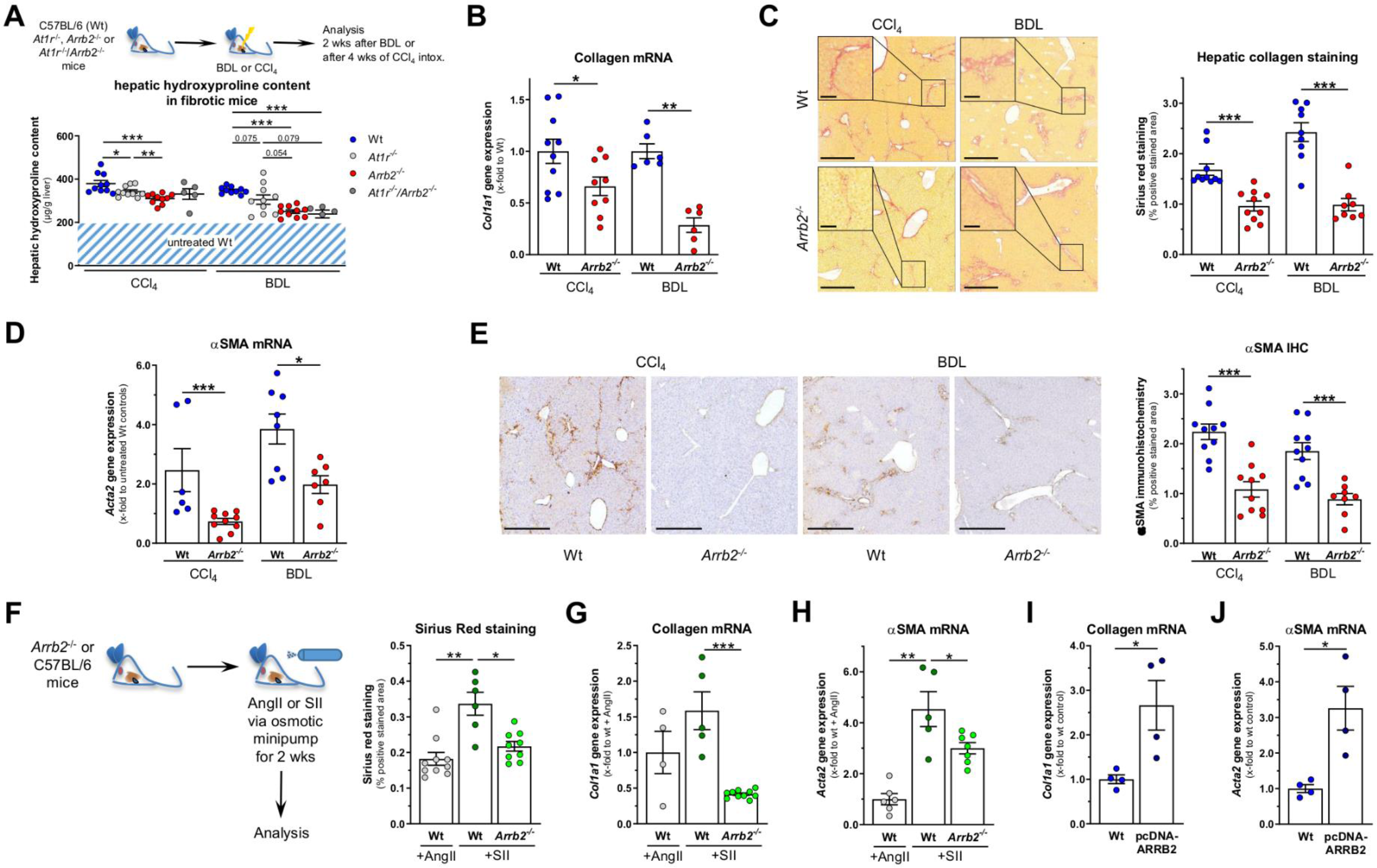
Inhibition and stimulation of beta-arrestin-2 in angiotensin II receptor type 1-mediated fibrogenesis. (A) Hepatic hydroxyproline content in wild-type (Wt), beta-arrestin-2 knock-out (*Arrb2-/-*), angiotensin II receptor type I A (AT1R) knock-out (*Agtr1-/-*) and *Arrb2-/-*/Agtr1-/- double knock-out mice in fibrosis models and in untreated Wt mice. Fibrosis was induced by carbon tetrachloride (CCl_4_) intoxication or by bile duct ligation (BDL). (B) Hepatic collagen 1 gene (*Col1a1*) expression in Wt and *Arrb2-/-* mice after CCl_4_ or BDL. (C) Representative Sirius red stainings of Wt and *Arrb2-/-* mice after CCl_4_ or BDL with magnification of fibrotic areas and the respective quantifications. Scale bars are 500µm and 100µm for the magnifications. (D) Hepatic alpha-smooth muscle actin (αSMA) gene (*Acta2*) expression in Wt and *Arrb2-/-* mice after CCl_4_ or BDL. (E) Representative sections of αSMA immunohistochemistry from Wt and *Arrb2-/-* mice after CCl_4_ or BDL and the respective quantifications. Scale bars are 500µm. (F) AT1R downstream signaling was stimulated G-protein-dependent by angiotensin II (AngII) or G-protein-independent via beta-arrestin-2 (ARRB2) by the peptide [Ser(1), Ile(4), Ile(8)]-angiotensin II (SII) for 14 days using osmotic mini-pumps. Quantification of Sirius red staining in Wt and Arrb2- /- mice after AT1R stimulation with AngII or SII. For representative stainings see Supp. Figure 1A. (G) Hepatic *Col1a1* expression in Wt and *Arrb2-/-* mice after AT1R stimulation with AngII or SII. (H) Hepatic *Acta2* expression in Wt and *Arrb2-/-* mice after AT1R stimulation with AngII or SII. (I) *Col1a1* expression in LX2 cells transfected with ARRB2 plasmids or control plasmids. (J) *ACTA2* expression in LX2 cells transfected with ARRB2 plasmids or control plasmids. Graphs show single measurements as dots (n=4-10 per group) and mean ± SEM. # = *p* 0.1 - 0.05, * = *p* < 0.05, ** = *p* < 0.01, *** = *p* < 0.001 by Mann-Whitney-U test.

Thereafter, the role of ARRB2 in HSC, the main producers of ECM in the liver (4), was investigated. HSC activation was measured by the surrogate marker alpha smooth muscle actin (αSMA) and its gene expression (*Acta2*). *Arrb2*-deficient mice showed significantly less HSC activation, as demonstrated by decreased *Acta2* expression (Figure 1D) and decreased αSMA positive staining (Figure 1E).

To further elucidate the impact of ARRB2 on HSC activation and fibrogenesis, experiments were performed using the full AT1R agonist angiotensin II (AngII) or a beta-arrestin-selective [Ser(1), Ile(4), Ile(8)]-angiotensin II (SII). The modified peptide SII selectively activates ARRB2 after binding to AT1R, while it is unable to activate Gα_q_-mediated signaling (12). In this experimental setting, wild-type (Wt) mice treated with AngII for two weeks via osmotic mini-pumps acted as controls. Stimulation of the beta-arrestin-specific pathways in Wt mice was performed by SII. In contrast, loss of function was achieved by stimulation with SII in *Arrb2*-deficient mice. Stimulation with SII in Wt mice increased Sirius red staining (Figure 1F, Supp. Figure 1A) and collagen expression (Figure 1G), as well as HSC activation, as shown by expression of the surrogate marker *Acta2* (Figure 1H), while SII had no effect in *Arrb2* deficient mice. Taken together, these data demonstrate that ARRB2 mediates hepatic fibrosis downstream of AT1R. *ARRB2* overexpression in transfected LX-2 cells, a human hepatic stellate cell line, led to increased gene expression of collagen 1 and αSMA (Figure 1I, J and Supp. Figure 1B).

### Processes involved in beta-arrestin-2-mediated fibrogenesis

RNA sequencing data from 23 cirrhotic patients and 7 non-fibrotic individuals demonstrate that hepatic *ARRB2* gene expression is upregulated in fibrosis in comparison to non-fibrotic livers, demonstrating the relevance of *ARRB2* in human disease (Figure 2A). By analyzing the same dataset with respect to *ARRB2* mediated effects, we found that transcriptional levels of *ARRB2* strongly correlate with the genes of extracellular signal-regulated kinases 1/2 (*MAPK1* and *MAPK3*), as well as with genes that indicate an interaction with activated hepatic stellate cells (*COL1A1* and *ACTA2*). In addition, ARRB2 negatively correlates with albumin gene expression (*ALB*), which is a marker of hepatocyte function (Figure 2B). The findings in humans were confirmed in a murine model of CCl_4_-induced fibrosis (Figure 2C, Supp. Figure 1C), demonstrating the suitability of the applied animal models for analysis of these mechanisms. These findings are in line with previously published human data from other cohorts and BDL-induced experimental fibrosis (13).

**Fig. 2.**
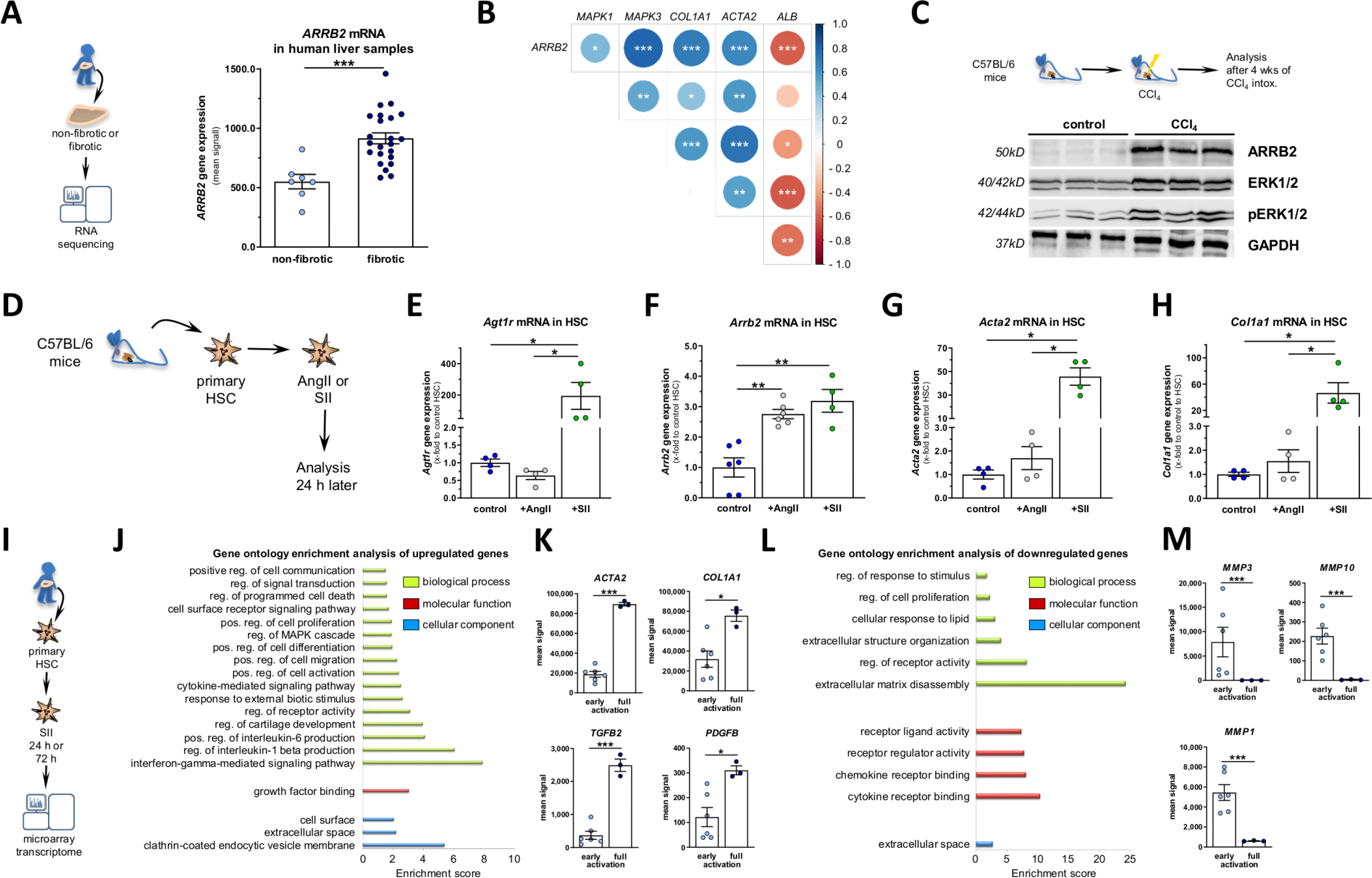
Processes involved in beta-arrestin-2-mediated fibrogenesis. (A) Hepatic beta-arrestin-2 (*ARRB2*) gene expression in human fibrotic and non-fibrotic liver tissue. (B) Correlation of *ARRB2* with markers of downstream signaling via extracellular-signal regulated kinase 1/2 (ERK1/2) (*MAPK1, MAPK3*), hepatic stellate cell (HSC) activity markers (*COL1A1, ACTA2*) and *ALB* for hepatocyte function in transcriptome data from whole liver tissue. (C) Representative Western blots (WB) of *ARRB2* and downstream signaling via ERK1/2 and its phosphorylated activated state. See quantification of WB in Supp. Figure 1C. (D) Primary HSC were isolated form Wt mice and stimulated G-protein-dependent by angiotensin II (AngII) or G-protein-independent via *ARRB2* by the peptide [Ser(1), Ile(4), Ile(8)]-angiotensin II (SII) for 24 hrs. (E) AT1R (*Agtr1*), (F) ARRB2 (*Arrb2*), (G) alpha-smooth muscle actin (αSMA; *Acta2*), (H) collagen 1 (*Col1a1*) gene expressions in murine primary HSC after AT1R stimulation with AngII or SII. (I) Experimental set-up for transcriptome analysis of primary human HSC harvested 24 hrs (early activation) and 72 hrs (full activation) after stimulation with SII. (J) Gene ontology enrichment analysis of upregulated genes comparing early with full activation showing biological processes, molecular functions and cellular components. (K) Representative upregulated genes comparing early and full activation with *ACTA2, COL1A1*, transforming growth factor-beta 2 (*TGFB2*) and platelet-derived growth factor subunit B (*PDGFB*). (L) Gene ontology enrichment analysis of downregulated genes comparing early with full activation showing biological processes, molecular functions and cellular components. (M) Representative downregulated genes comparing early and full activation with matrix metalloproteinases *MMP3, MMP10* and *MMP1*. Graphs show single measurements as dots (n=7-23 for whole liver samples and n=4-6 for primary cell culture) and mean ± SEM. * = *p* < 0.05, ** = *p* < 0.01, *** = *p* < 0.001 by Mann-Whitney-U test, Spearman correlation (for B) or false discovery rate (FDR) adjusted t-test (for K, M).

Since the results indicate a role of ARRB2 in HSC activation with HSC being the major drivers of hepatic fibrosis (4), stimulation of ARRB2 activity was investigated more closely in primary murine and primary human HSC. For this purpose, primary murine HSC were incubated *in vitro* for 24 hrs with ARRB2-specific agonist SII or the unspecific peptide AngII (Figure 2D). Incubation with SII significantly increased expression of *Agtr1* and *Arrb2* as well as markers for HSC activation (*Acta2*) and fibrosis (*Col1a1*), while incubation with AngII led only to upregulation of *Arrb2* and to non-significant mild effects on *Agtr1*, *Acta2* and *Col1a1* expressions (Figure 2E-H).

Transcriptome analyses were performed by RNA microarray in primary human HSC incubated with SII for 24 hrs or 72 hrs to elucidate time-dependent effects of ARRB2 stimulation at the stage of early and full HSC activation (Figure 2I; Supp. Figure 1D). Although HSC are activating over time in culture, incubation with SII for 72 h had an even stronger effect on *RARRES1*, *ACTA2* and *COL1A1* gene expression than unstimulated auto-activation of the HSC during 72 h in culture (Supp. Figure 1E). Gene set enrichment analysis clusters sets of genes that are associated to a specific biological process, molecular function or cellular component. Analysis of upregulated genes revealed processes associated with HSC activation, such as proliferation, migration and differentiation, along with proinflammatory processes and cell communication (2)(Figure 2J; Supplemental Table 1). Among the cellular components that were affected, clathrin-coated endocytic vesicle membrane showed the highest enrichment (Figure 2J). ARRB2 is involved in endocytosis and desensitizes AT1R via this mechanism (14). Single genes that were upregulated in samples incubated with SII for 72 hrs were related to HSC activation (*ACTA2*, *TGFB2*, and *PDGFB*) and fibrosis (*COL1A1*) (Figure 2K) compared to samples incubated for 24 hrs. For downregulated genes, incubation with SII for 72 hrs affected mainly the extracellular matrix and especially its disassembly (Figure 2L; Supplemental Table 2). Thus, matrix metallopeptidases were among the single genes that were significantly downregulated after 72 hrs of incubation with SII (Figure 2M).

These data strongly support the hypothesis that ARRB2 is a mediator of HSC activation and crucially involved in key processes of fibrosis.

### Interaction of beta-arrestin-2 and retinoic acid receptor responder 1 in fibrosis

ARRB2-specific stimulation by SII led to upregulation of genes responsible for HSC activation and fibrosis formation. Hence, in order to identify major regulators of these processes, correlation analyses with key markers of HSC activation were performed for potential candidates, which were pre-selected by literature research. One of them is retinoid acid receptor responder 1 (*RARRES1)*, which has been described as a shared regulator in liver, kidney and lung fibrosis (3), and was among the upregulated genes after stimulation of ARRB2-specific signaling by SII (Figure 3A). Furthermore, *RARRES1* strongly correlated with markers of HSC activation (*ACTA2*, *PDGFB*, *TGFBR1*, *TGFB2*), inflammation (*CCL2*) and accumulation of ECM (*COLA1A1*, *COL4A1*, *FN1*), while it negatively correlated with ECM degradation markers (*MMP9*) (Figure 3A).

**Fig. 3.**
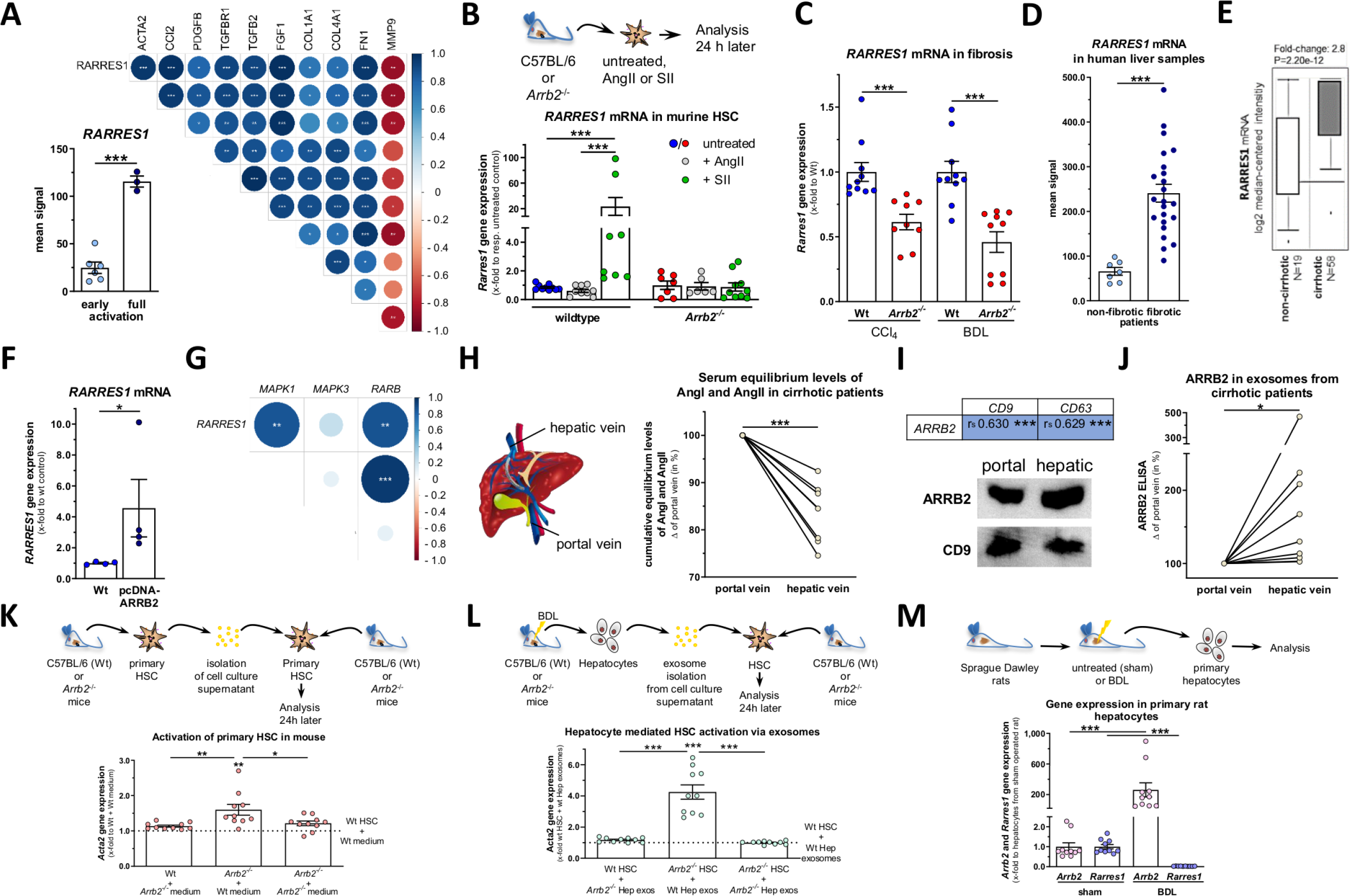
Role of RARRES1 in fibrosis. (A) Retinoic acid receptor responder protein 1 (*RARRES1*) expression in primary human hepatic stellate cells (HSC) harvested 24 hrs (early activation) and 72 hrs (full activation) after stimulation with [Ser(1), Ile(4), Ile(8)]- angiotensin II (SII) and correlation matrix of *RARRES1* and markers of HSC activation (*ACTA2, CCL2, PDGFB, TGFBR1, TGFB2* and *FGF1*), collagen deposition (*COL1A1, COL4A1* and *FN1*) and degradation (*MMP9*). (B) *Rarres1* expression in primary HSC isolated from wild-type (Wt) and beta-arrestin-2 knock-out (*Arrb2-/-*) mice after AT1R stimulation with AngII or SII. (C) *RARRES1* expression in whole liver samples from Wt or *Arrb2-/-* mice after carbon tetrachloride (CCl_4_) intoxication or bile duct ligation (BDL). (D) *RARRES1* expression in human fibrotic and non-fibrotic liver tissue. (E) *RARRES1* expression in an external validation cohort (15). (F) *RARRES1* expression in LX2 cells transfected with *ARRB2* plasmids or control plasmids. (G) Correlation of *RARRES1* with markers of downstream signaling via extracellular-signal regulated kinase 1/2 (*MAPK1, MAPK3*) and retinoic acid receptor beta (*RARB*). (H) Cumulative serum equilibrium levels of AngI and AngII in portal and hepatic vein samples from cirrhotic patients. (I) Spearman correlation (r_s_) of *ARRB2* with exosome markers *CD9* and *CD63* in whole liver transcriptome data, as well as protein levels of ARRB2 and CD9 and (J) quantification of circulating *ARRB2* levels in portal and hepatic vein exosomes from cirrhotic patients. (K) *Acta2* expression in primary HSC isolated from Wt or *Arrb2-/-* mice and cell culture supernatant for measurement of HSC-HSC communication. (J) *Acta2* expression in primary mouse HSC and injured hepatocytes after BDL isolated from Wt or *Arrb2-/-* mice and exosomes isolated from cell culture supernatant to assess hepatocyte-HSC communication. (K) Arrb2 and *RARRES1* expression in rat primary hepatocytes from healthy sham-operated or fibrotic BDL rats. Graphs show single measurements as dots (n=4-10 except for transcriptome data of hHSC with n=3-6 and whole liver transcriptome data with n=7-23) and mean ± SEM or bar graphs with mean ± SEM and *n* value given for each group. * = *p* < 0.05, ** = *p* < 0.01, *** = *p* < 0.001 by Mann-Whitney-U test, Spearman correlation (for A, G, I) or false discovery rate (FDR) adjusted *t*-test (for A).

To verify whether *RARRES1* upregulation is ARRB2-dependent, primary HSC from wild-type and *Arrb2*-deficient mice were incubated with AngII or SII. While AngII did not upregulate *Rarres1* expression in Wt HSC, incubation with SII resulted in a marked upregulation. In HSC from *Arrb2*-deficient mice, none of the stimulants led to upregulation of *Rarres1* (Figure 3B). Furthermore, *Rarres1* expression was diminished in two different fibrosis mouse models with *Arrb2* deficiency (Figure 3C).

Next, the relevance of RARRES1 to human disease was investigated. Upregulation of *RARRES1* gene expression was demonstrated not only by *in vitro* stimulation of human HSC with SII, but also in liver samples from patients with fibrosis (Figure 3D). This was confirmed *in silico* in an external cohort of 77 patients, 58 of whom were patients with cirrhosis (15) (Figure 3E). Transcriptome analysis revealed strong positive correlation of *RARRES1* with *COL1A1* and *ACTA2* and negative correlation with *ALB* in human liver samples (Supp. Figure 2A), similar to those correlations presented in Figure 2 for *ARRB2*, highlighting the potential involvement of RARRES1 in fibrogenesis and suggesting an association with hepatocyte injury marked by decreased hepatocyte function. The dependency on ARRB2 for *RARRES1* gene expression was further endorsed by transfection of LX-2 cells with an ARRB2 plasmid, which significantly induced the gene expression of *RARRES1* in transfected cells (Figure 3F and Supp. Figure 1B). Besides many other functions, ARRB2 modulates the activity of the nuclear retinoic acid receptors (RAR) via ERK2 (*MAPK1*), with strongest effects on RARβ (16). It has been described in skin grafts that RARRES1 is one of the retinoic acid responsive genes and is regulated by RARβ and RARγ (17). Transcriptome data from primary human HSC as well as whole liver tissue indicate that *RARRES1* expression in fibrogenesis is dependent on this pathway via ARRB2 and downstream signalling via ERK2 and RARβ (Figure 3G and Supp. Figure 2B).

Variations in *Arrb2* expression during different stages of HSC activation were observed in freshly isolated and culture activated HSC. Freshly isolated cells from fibrotic BDL rats showed higher *Arrb2* expression than cells isolated from sham-operated rats and activated in vitro (Supp. Figure 2C), which usually is a far stronger stimulus (18). Thus, we hypothesized that ARRB2 in vivo may also be transferred from other sources. Levels of AngII and ARRB2 were measured in portal vein and hepatic vein blood samples from cirrhotic patients collected during transjugular portosystemic shunt insertion, a catheter-based technique accessing the portal vein (19). Serum equilibrium levels of angiotensin I (AngI) and AngII in the portal vein were significantly upregulated, probably due to higher absorption of angiotensinogen and renin in the liver, resulting in enhanced local formation of AngII in the liver (Figure 3H; Supp. Figure 2E). To elucidate the major source of ARRB2, exosomes in the blood samples from these two vascular beds were investigated. Pironti *et al.* showed that exosomes may transport AGTR1 from one organ to another (20). It is known that ARRB2 is attached to AT1R for its internalization and vesicle formation. We could demonstrate a significant upregulation of the exosome markers *CD9* and *CD63* in fibrotic livers (Supp. Figure 2D) and a strong correlation with *ARRB2* at mRNA level (Figure 3I). Furthermore, we detected ARRB2 and CD9 protein in exosomes isolated from portal and hepatic vein samples (Figure 3I). ARRB2 protein, measured by ELISA, mainly derives from the liver as demonstrated by significantly higher levels of ARRB2 in exosomes from the hepatic vein (Figure 3J).

Thus, the question arose as to whether ARRB2-loaded exosomes derive from activated HSC or other liver cells. As we could exclude any involvement of liver sinusoidal endothelial cells (Supp. Figure 3A), immune and Kupffer cells (Supp. Figure 3B-C), even after bone marrow transplantation and larger microvesicles (Supp. Figure 3D) for transport in ARRB2-mediated fibrosis, we focused on the interaction between hepatocytes and HSC. Primary HSC or hepatocytes were isolated from either Wt or *Arrb2*-deficient mice and were separately cultured for 24 hrs. Cell culture supernatant from these cultures was transferred to primary HSC cultures. HSC isolated from *Arrb2*-deficient mice could be activated by supernatant isolated from HSC of Wt mice, as demonstrated by increased *Acta2* expression (Figure 3K). Interestingly, *MIR119A*, which has been described as an inhibitor of HSC activation in HSC-HSC-communication (21), was downregulated, while *CTGF*, a stimulating factor for activation in HSC-HSC-communication (22), was upregulated in HSC treated with SII for 72h (Supp. Fig. 3E).

However, the effect on HSC activation was much more pronounced when HSC isolated from *Arrb2*-deficient mice were incubated with exosomes purified from cell culture supernatant of hepatocytes isolated from Wt mice (Figure 3L). In both experiments, none of the other possible combinations increased HSC activation (Figure 3K-L). Consequently, expressions of *Arrb2* and *Rarres1* were also investigated in hepatocytes isolated from either sham-operated or cirrhotic rats. While *Arrb2* was massively upregulated in hepatocytes from fibrotic animals, *Rarres1* was significantly downregulated in these cells (Figure 3M). These experiments suggest an interaction between hepatocytes and HSC with ARRB2 as a potential driver of intercellular communication via exosomes. The role of these exosomes was also investigated in the extrahepatic compartment, however, without significant effects on contractility of vessels (Supp. Figure 4). Therefore, this study demonstrates that ARRB2 induces fibrosis via RARRES1 induction and that ARRB2 is released from injured hepatocytes, possibly transferring the protein into HSC to maintain their fibrogenic phenotype.

**Fig. 4.**
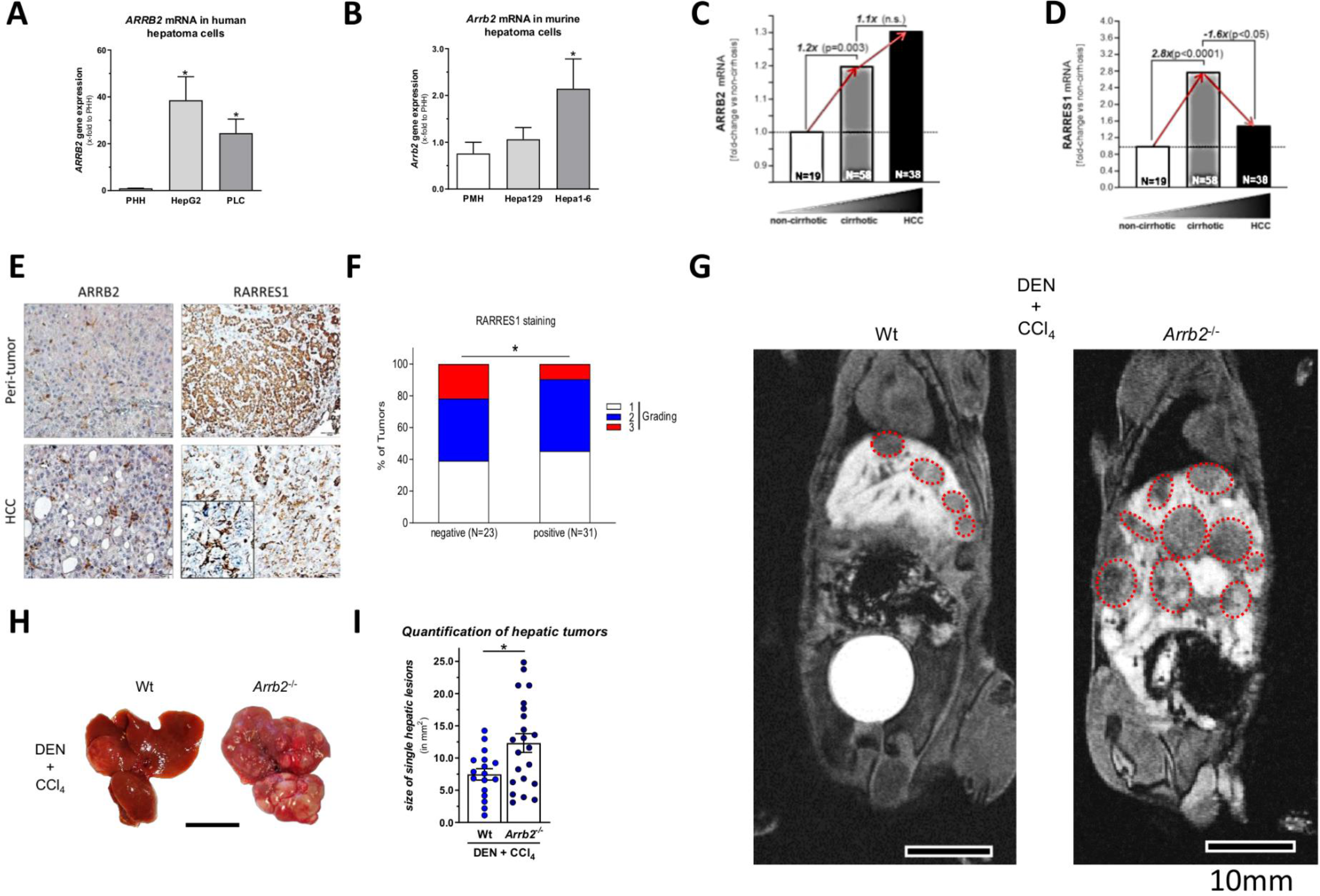
Role of β-arrestin-2 and RARRES1 in HCC. (A) *ARRB2* expression in primary human hepatocytes (PHH) and the hepatoma cell lines HepG2 and PLC. (B) *Arrb2* expression in primary mouse hepatocytes (PMH) and the hepatoma cell lines Hepa129 and Hepa1-6. (C) *In silico* analysis of *ARRB2* expression in liver tissue of patients with and without cirrhosis or with hepatocellular carcinoma (HCC). (D) *In silico* analysis of *RARRES1* expression in liver tissue of patients with and without cirrhosis or with HCC. (E) Representative sections of *ARRB2* and *RARRES1* immunohistochemistry in peri-tumor and HCC tissue micro arrays. (F) Semi-quantitative measurement of RARRES1 immunohistochemistry positive staining in tumor tissue in relation to tumor grades. (G) Magnetic resonance imaging of the liver in wild-type (Wt) or beta-arrestin-2 knock-out (*Arrb2*^-/-^) with a fibrosis and tumor generating model of single Diethylnitrosamine (DEN) injection and repetitive carbon tetrachloride (CCl_4_) intoxication. Hepatic lesions are highlighted with a red dotted line. Scale bars are 10 mm. (H) Morphological comparison of livers from Wt and *Arrb2*^-/-^ mice after single DEN injection and repetitive CCl_4_ intoxication. Scale bar is 10 mm. (I) Mean size of hepatic tumors in mice receiving a single DEN injection and repetitive CCl_4_ intoxication quantified in magnetic resonance imaging sections. Graphs show single measurements as dots and mean ± SEM or bar graphs with mean ± SEM and *n* value given for each group. * = *p* < 0.05, ** = *p* < 0.01 by Mann-Whitney-U test.

### Beta-arrestin-2/retinoic acid receptor responder 1 axis in the development of HCC

Since RARRES1 is a well-described tumor suppressor, interaction of RARRES1 with ARRB2 was investigated in the development of hepatocellular carcinoma (HCC) in fibrosis. *ARRB2* expression is upregulated in human (Figure 4A) and murine (Figure 4B) HCC cell lines compared to primary hepatocytes. Accordingly, *Rarres1* expression is decreased in human and murine liver cancer cell lines (Supp. Figure 5A-B). In addition to the increase of *ARRB2* expression in cirrhosis, a further, however non-significant, increase was measured in liver samples from patients with HCC (Figure 4C). The increase of ARRB2 expression in HCC was confirmed in murine liver tissue (Supp. Figure 5C). In contrast, *RARRES1* expression increased in cirrhosis in a similar way to *ARRB2* expression but was significantly attenuated in HCC tissue (Figure 4D). A slight, however not significant, decrease of RARRES1 expression could also be observed in murine HCC (Supp. Figure 5D), probably due to expression in non-tumorous cells. Furthermore, ARRB2 and RARRES1 were analyzed in human peri-tumor and HCC tissues applying a human tissue micro array (23). ARRB2 was detected in peri-tumorous hepatocytes and in peri-tumorous and stromal hepatic stellate cells, while within HCC tissues, RARRES1 was detected almost exclusively in hepatic stellate cells (Figure 4E) and RARRES1 positive staining was inversely correlated with tumor grading in HCC (Figure 4F). Additionally we used a fibrosis and tumor generating model of single Diethylnitrosamine (DEN) injection and repetitive CCl_4_ intoxication for 14 weeks in Wt and *Arrb2*-deficient mice. The number and size of tumors (red shapes) was dramatically increased in *Arrb2*-deficient mice compared to their Wt littermates as assessed by magnetic resonance imaging (MRI) and macroscopic observations of the harvested livers (Figure 4G-I). The degree of hepatic fibrosis was slightly decreased in *Arrb2*-deficient mice as assessed by Sirius red staining (Supp. Figure 5E).

**Fig. 5.**
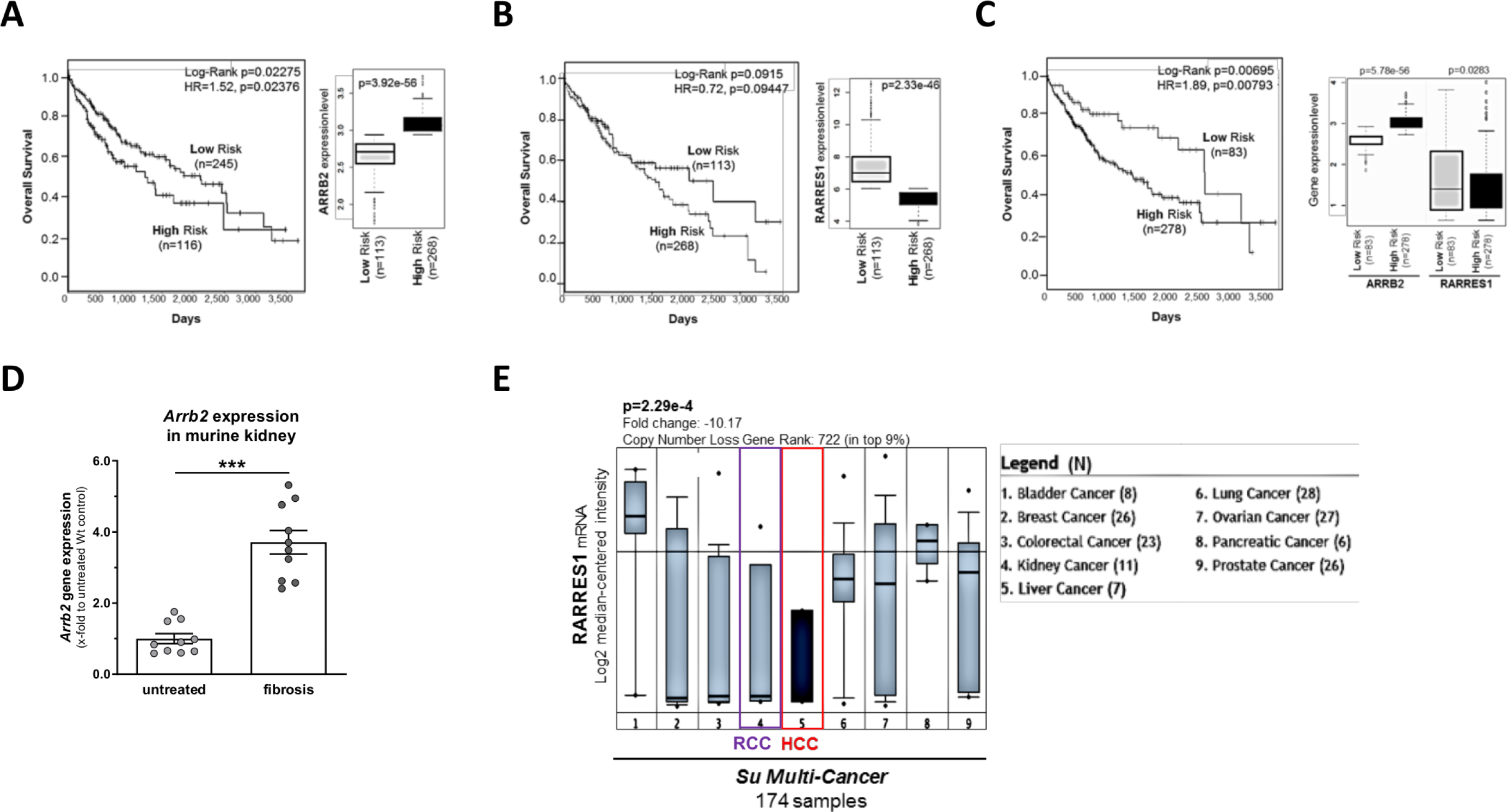
Role of beta-arrestin-2 and RARRES1 in cancer survival. (A) Survival curves of patients with HCC stratified by *ARRB2* expression profile. High *ARRB2* expression represents the high risk sub-cohort. (B) Survival curves of patients with HCC stratified by *RARRES1* expression profile. Low *RARRES1* expression represents the high risk sub-cohort. (C) Survival curves of patients with HCC stratified by combined *ARRB2*/*RARRES1* expression profile. High *ARRB2* expression and low *RARRES1* expression represents the high risk sub-cohort. (D) Quantification of *Arrb2* expression in murine kidney fibrosis using the adenine-induced tubulointerstitial nephritis model compared to control samples from healthy animals. (E) *In silico* analysis of *RARRES1* expression in various human organ cancers. HCC is highlighted by a red frame and kidney cancer (RCC) by a purple frame. Graphs show single measurements as dots (n=10) and mean ± SEM or bar graphs with mean ± SEM and *n* value given for each group. The number of patients used for survival curves and *in silico* analysis is indicated in each panel. *** = *p* < 0.001 by Mann-Whitney-U test.

### Role of beta-arrestin-2 and retinoic acid receptor responder 1 in cancer survival

Since the beta-arrestin-2/retinoic acid receptor responder 1 axis seems to play an important role in HCC, its impact on survival was investigated. Analysis of The Cancer Genome Atlas (TCGA)-derived patient datasets applying the SurvExpress Database revealed enhanced *ARRB2* and reduced *RARRES1* expression in high-risk compared with low-risk (computational stratification was based on prognostic index) HCC patient groups. Moreover, enhanced *ARRB2* (Figure 5A) and reduced *RARRES1* (Figure 5B) expression levels were associated with poor overall survival in HCC patients. Indeed, combined enhanced *ARRB2* and reduced *RARRES1* levels in high-risk patients revealed synergistic effects on overall patient survival (Figure 5C).

Since *ARRB2* is ubiquitously expressed, the demonstrated *ARRB2*/*RARRES1* interaction could also play an important role in fibrotic processes and the development of cancer in other organs. We thus investigated the expression of both mRNAs in fibrotic kidneys and in diverse cancer types including liver and kidney cancer. Similar to the liver, *Arrb2* was upregulated in murine kidney fibrosis in a model of adenine-induced tubulointerstitial nephritis (Figure 5D). Furthermore, *RARRES1* expression was evaluated in *in silico* diverse cancer tissues and found to be downregulated when compared to average expression levels in cancer tissues in, e.g., HCC. It also showed low levels in kidney cancer (RCC) (Figure 5E). Thus, targeting or interfering of the ARRB2/RARRES1 axis may prevent organ fibrosis. Furthermore, expression levels and ratios could be used as biomarkers to determine the risk to develop cancer after fibrosis in multiple organs.

## Discussion

This study demonstrates the important role of beta-arrestin-2 (ARRB2) in liver fibrogenesis and cellular crosstalk, elucidating the molecular basis for transition from fibrosis towards malignancy. Myofibroblasts transdifferentiated from activated HSC are the major contributors to liver fibrogenesis due to extensive synthesis and release of ECM (4). In this study, selective stimulation of AGTR1 in HSC increased ARRB2. As a result, HSC activation markers and collagen synthesis were increased. AngII, the agonist of AT1R, is known to induce proliferation and contractility in HSC (24, 25). In previous studies, the important role of G protein-dependent signaling downstream of AT1R was described in liver fibrosis and portal hypertension (5, 6). However, studies have also shown that AT1R regulates G protein-independent signaling and recent findings suggest that ARRB2 mediates increased proliferation in HSC, which is also G protein-independent, via ERK1/2 (26, 27). These two pathways downstream of AT1R, G protein-dependent and G protein-independent, are probably meant to maintain cellular homeostasis and therefore trigger different mechanisms. This study emphasizes the importance of ARRB2-dependent signaling in HSC activation and liver fibrogenesis.

In experimental and human liver fibrosis, ARRB2 and ERK1/2 activity were increased, while *Arrb2* deficiency led to decreased liver fibrosis, probably mediated by decreased HSC activation. As shown by Li et al. and Lovgren et al., the function of ARRB2 in HSC could be similar to its function in cardiac and pulmonary fibroblasts (8, 28). Both studies emphasize the important role of ARRB2 in fibroblasts for development of fibrosis and collagen synthesis in the respective organs. In liver fibrosis, this function is assumed by activated HSC, which transdifferentiate into a myofibroblast-like phenotype (4). Additionally, our data suggest a role for ARRB2 in kidney fibrosis. These studies also show that, depending on the organ, treatment of fibrosis by inhibition of ARRB2 needs to be organ-specific, and preferably cell-specific. At least for the liver, such carriers, which target HSC exclusively and onto which an ARRB2 inhibitor could be bound, exist (29, 30). Also, anti-*ARRB2* RNA aptamers coupled to a cancer cell-specific carrier have been described to inhibit ARRB2 activity (31). Furthermore, deficiency of *Arrb2* was shown to be more effective in decreasing fibrogenesis than deficiency of upstream *Agtr1*, making ARRB2 a potent target for antifibrotic treatment. In contrast to ARRB2, ARRB1 remains unchanged during progressive liver injury, indicating that ARRB1 is not involved in HSC activation and liver fibrogenesis (27).

Upregulation of *RARRES1* has been described as playing a pivotal role in liver fibrosis (3). However, while mechanisms by which RARRES1 is regulated were lacking, this study clearly demonstrates the close relationship of ARRB2 and RARRES1 in fibrogenesis. Deficiency of *Arrb2* precipitates downregulation of *Rarres1* in experimental models of liver fibrosis, resulting in less collagen accumulation. Since it has been shown previously that *Rarres1* is already upregulated in the early stages of HSC activation (3), it seems to be obvious that ARRB2 mediates HSC activation and, consequently, fibrogenesis via RARRES1. Thereby, ARRB2 might regulate RARRES1 expression via ERK2 and the nuclear RARβ (16, 17).

Apart from HSC, *Arrb2* and *Rarres1* are also expressed in other hepatic cell types. Interestingly, *Arrb2* was found to be upregulated in hepatocytes upon fibrosis induction, while *Rarres1* was downregulated in hepatocytes of fibrotic livers. Nevertheless, absence of *Arrb2* has been shown to improve hepatocyte and overall survival in experimental cholestatic liver fibrosis in mice. These mechanisms are mediated by activation of Akt and downregulation of its target GSK3β (32). However, increased hepatocyte survival in the absence of *Arrb2* cannot explain decreased fibrosis upon liver injury. Therefore, another finding of this study may be of great importance, namely that we could demonstrate that exosomes released by hepatocytes were able to increase RARRES1 and αSMA even in the absence of *Arrb2*. ARRB2, and possibly RARRES1, which has been detected in exosomes from the parotid gland (33) and which are transferred from hepatocytes to HSC, mediate these effects. This finding indicates a distinct hepatocyte/HSC communication leading to activation of HSC.

In cirrhotic patients, serum equilibrium levels of AngII were higher in the pre-hepatic portal vein than in the post-hepatic vein, and vice versa for ARRB2. Given that serum equilibrium levels of AngII are directly affected by the concentration of renin and angiotensinogen, we hypothesized that these factors entering the liver via the portal vein are absorbed, leading to local interstitial formation of AngII and stimulating hepatic cells to synthesize and release ARRB2. Pironti et al. recently reported that biomechanically stressed cardiomyocytes release AT1R-containing exosomes, which target other cardiomyocytes, skeletal myocytes and mesenteric vessels. However, the exosomes contained AT1R only in the presence of ARRB2 (20).

In the hepatic outflow, high levels of circulating ARRB2 could also account for the hypocontractility of vessels in liver cirrhosis. One possible theory is that ARRB2-containing exosomes are absorbed by the smooth muscle cells of the vessels leading to desensitization of the AT1 receptor and decreased Ca^2+^ sensitivity (34). As in cardiomyocytes in the heart, the origin of hepatic exosomes containing ARRB2 could be activated HSC with their myofibroblast-like phenotype or LSEC with their response mechanisms to blood flow-induced biomechanical stress. However, while we could exclude any influence of these exosomes on extrahepatic vasodilation, a possible hypothesis is that ARRB2 exosomes are meant to be delivered to LSECs, which lose ARRB2 upon injury, a process that seems to be related to endothelial dysfunction, as demonstrated recently (35). In fibrosis, the exosomes do not pass the space of Disse to reach the LSECs but are instead taken up by HSC to maintain their profibrotic phenotype.

Due to its characteristic as a tumor suppressor, we also investigated ARRB2/RARRES1 interaction in HCC. In contrast to its close relationship to fibrosis, RARRES1 is uncoupled from ARRB2 signaling in HCC. The downregulation of *RARRES1* in HCC and other cancers is mediated by hypermethylation of *RARRES1* (36, 37), which seems to be independent of ARRB2. Since RARRES1 improves survival in HCC, inhibition of hypermethylation potentially halts HCC progression.

In conclusion, the present study found that ARRB2 is upregulated upon AT1R stimulation leading to increased HSC activation and collagen synthesis mediated by RARRES1 downstream of ARRB2. In liver fibrosis, ARRB2-enriched exosomes released by hepatocytes promote HSC activity and their transdifferentiation into a myofibroblast-like phenotype. Decreased collagen accumulation in *Arrb2* deficiency contributes to diminished HSC activation by decreasing *Rarres1* expression. In HCC, RARRES1 is uncoupled from ARRB2 signaling. Furthermore, ARRB2 released from fibrotic livers is probably AngII-mediated and may contribute to hypocontractility in extra-hepatic vessels. Therefore, this study unveils novel processes and mechanisms involved in HSC activation and the consequences thereof for liver fibrosis and cancer. Importantly, we identified the intercellular communication of extracellular vesicles contributing to intra- and extrahepatic effects.

## Materials and methods

### Human samples

Liver samples of cirrhotic patients were taken during liver transplantation. Non-cirrhotic samples were taken from patients undergoing liver resection. Following specimen collection, samples were snap-frozen. None of the donors received angiotensin-converting enzyme inhibitors, angiotensin receptor agonists or catecholamines before surgery (5, 6, 38). Blood samples from the portal vein and the hepatic vein were collected during insertion of transjugular intrahepatic portosystemic shunt (TIPS) as described previously (39, 40).

### Animals

For animal studies, male wild-type (Wt, C57BL6/6J background; Charles River), beta-arrestin-2 knock-out (*Arrb2*^-/-^, C57BL/6J background), angiotensin II receptor type I subtype A knock-out (*Agtr1*^-/-^, C57BL/6J background) and *Arrb2*^-/-^/ *Agtr1*^-/-^ double knock-out mice were used. *Agtr1*^-/-^ were kindly provided by Nikos Werner (Department of Internal Medicine II, University of Bonn, Germany) and *Arrb2^-^*^/-^ mice were kindly provided by Robert Lefkowitz (Department of Medicine, Duke University Medical Center, USA). We only investigated the knock-out of the subtype A of the AT1R, since subtype B is not expressed in murine livers (41). Furthermore, Sprague-Dawley wild-type (WT) rats were used for hepatic cell isolation. All animals were kept at 22°C with a 12 hrs day/night cycle and received water and chow *ad libitum*.

### Cholestatic model of fibrosis

Bile duct ligation (BDL) was performed in adult mice with an initial weight of 20-25g as described previously (5, 42). Sham-operated mice served as control. Experiments were performed 14 days after BDL when animals developed severe hepatic fibrosis.

### Toxic model of fibrosis

Periodic carbon tetrachloride (CCl_4_) inhalation of 1L/min was performed thrice weekly in adult mice with an initial body weight of 20-25g as described previously (5, 42, 43). To enhance hepatotoxicity, mice were treated with 0.3g/L phenobarbital in drinking water during the time of fibrosis induction. Experiments were performed after four weeks of treatment when the animals developed severe hepatic fibrosis.

### Model of hepatocellular carcinoma with fibrosis

Diethylnitrosamine (DEN; 1mg/kg) was injected i.p. in 14 day old Wt or *Arrb2*^-/-^ mice. Starting at an age of 8 weeks mice were administered CCl_4_ twice a week for 14 weeks as described elsewhere (44). Corn oil (Sigma Aldrich, Germany) was used as solvent. Control animals received only solvent. Experiments were performed after 14 weeks of CCl_4_ at an age of 22 weeks.

### Adenine-induced tubulointerstitial nephritis

The adenine-induced tubulointerstitial nephritis model was used to cause chronic inflammation and end-stage fibrosis as described previously (45). Eight to 12 week old C57BL6/6J were fed with an adenine-enriched diet (Sniff, Soest, Deutschland) for 21 days as described elsewhere using a lower adenine concentration of 0.2% (45, 46).

### HCC visualization by magnetic resonance imaging

Size and number of hepatic tumors was investigated in anesthetized mice with contrast-enhanced magnetic resonance imaging (MRI) using gadoxetic acid (Gd-EOB-DTPA, 0.057 mmol/kg, Primovist, Bayer-Schering, Berlin, Germany) as contrast agent and was performed on a 3T MRI scanner (Siemens Magnetom Trio, Siemens Medical Solutions). The animals were imaged in the axial plane and the imaging sequences included T1-weighted 3D gradient echo sequences with an echo time of 20 milliseconds, a repetition time of 947 milliseconds, a field of view of 100 mm, a slice thickness of 1 mm, a flip angle of 140° and fat suppression for saturated fat as described elsewhere (47, 48). Quantification of hepatic tumors was performed by ImageJ (version 1.51q, NIH, USA).

### Tissue collection

Before animals were sacrificed after completion of the experiments, they were fasted overnight with free access to water. Animals were sacrificed by cervical dislocation for tissue collection. Liver specimens were snap-frozen in liquid nitrogen and stored at -80°C for molecular biologic experiments or fixed in formaldehyde (4%) for histological staining as described previously (49).

### Hepatic hydroxyproline content

The content of collagen was determined by the photometrical measurement of the major component hydroxyproline in liver hydrolysates using standard protocols (49).

### Isolation of primary human HSC

Primary human HSC (hHSC) were kindly provided by Krista Rombouts (University College London, UK) and isolated as described previously (50). Experiments described in this study were performed with hHSC cultured from at least three cell preparations (donors) and three replicates, used between passage 6 and 9.

### Isolation of human liver sinusoidal endothelial cells

Human cells were isolated from residue tissue obtained after liver transplantation and after partial hepatectomy to excise tumor metastasis from colon carcinoma, while only healthy cells from peritumoral tissue confirmed as “normal” by anatomical pathologists were used as controls from the latter tissue. Ethics Committee of the Hospital Clínic de Barcelona approved the experimental protocol (HCB/2015/0624), and patients signed an informed consent. LSEC were isolated and cultured using standardized protocols as described previously (51).

### Isolation of primary liver cells from mice and rats

Murine primary hepatocytes and hepatic stellate cells were isolated from male Wt, or *Arrb2*^-/-^ mice following standard protocols as described previously (38, 52). Rat primary hepatocytes and hepatic stellate cells were isolated from male WT rats as described previously (5, 52). Experiments were performed either in freshly isolated cells (1-7 days after isolation) or culture activated cells (2. And 3. passage).

### Calcium phosphate transfection

ARRB2_OHu23516C_pcDNA3.1(+)-C-eGFP and pcDNA3.1(+)-C-eGFP control plasmids were purchased from Genescript (Piscataway, USA). LX2 cells were seeded on 6-well plates (300,000 cells/well) in DMEM medium (10% FBS, 1% Penicilin/Streptomycin and 2% L-Glutamine) and were grown to 80% confluency. For cell transfection, BBS buffer (50 mM BES, 280 mM NaCl and 1.5 mM Na2HPO4) was preprared in sterile water and pH was adjusted to 6.96. Before transfection, the medium was replaced with serum free medium (without FBS). The transfection mix was prepared by adding sterile water, the corresponding plasmid, BBS buffer and 2 M CaCl_2_. The transfection mix was added drop by drop in each well, and cells were incubated for 2.5 h. Afterwards, the medium was replaced with DMEM medium and cells were incubated for 48 h before harvest for further analysis.

### Generation of bone marrow chimera, depletion of hepatic macrophages and macrophage transfer

Bone marrow chimeric animals were generated by irradiating mice with 9Gy and reconstitution with 5 × 10^6^ donor bone marrow cells injected into the tail vein. To adoptively transfer bone-marrow-derived macrophages and neutrophils a MACS isolation procedure was used as described previously (53). 30μl of clodronate or containing liposomes or empty liposomes as control (Encapsula) were injected in the foot-pad of mice before macrophage transfer as described previously (54, 55). Clodronate injection did not induce granuloma formation at the site of injection nor did it enlarge the depleted lymph node.

### Human and murine hepatocellular carcinoma cell lines

Primary human hepatocytes were cultured as described previously (23, 56). Additionally, human-derived commercially available HCC cell lines PLC (ATCC CRL-8024, LGC Standards, Wesel, Germany) and HepG2 (ATCC HB-8065, LGC Standards, Wesel, Germany) were cultured as described by others (23, 57). Furthermore, murine HCC cell lines Hepa129 (NCI-Frederick Cancer Research and Development Center, DCT Tumour Repository) and Hepa1-6 (ATCC CRL-1830) were used and cultured as described previously (23).

### Organ chamber experiments

Circulating exosomes from mice were pelleted as described previously and resuspended in Dulbecco’s modified Eagle medium (DMEM) filtered on 0.1 μm membrane pores to reach a final volume corresponding to the initial volume of plasma. The effect of exosomes was compared with the effect of the same volume of filtered DMEM (58, 59).

Thoracic aortas from adult C57BL/6 mice (eight to ten weeks old) were isolated after animal sacrifice under isoflurane anesthesia. Mouse aortic rings were incubated for 24 hrs; 37°C in a 5% CO_2_ incubator with filtered DMEM supplemented with antibiotics (100 IU/mL streptomycin, 100 IU/mL penicillin [Gibco, Invitrogen, Paisley, Scotland], and 10 μg/mL polymyxin B [Sigma, St Louis, USA]) either alone or in the presence of circulating exosome from Wt or *Arrb2*^-/-^ mice. After this incubation period, the rings were mounted in organ chambers (Multi Wire Myograph System, model 610 M; Danish Myo Technology, Aarhus, Denmark) filled with Krebs–Ringer solution (NaCl 118.3 mmol/L, KCl 4.7 mmol/L, MgSO_4_ 1.2 mmol/L, KH_2_PO_4_ 1.2 mmol/L, CaCl_2_ 1.25 mmol/L, NaHCO3 25.0 mmol/L and glucose 5.0 mmol/L) gassed with a mixture of O_2_ 95% and CO_2_ 5% (pH 7.4). The presence of functional endothelial cells was confirmed by the relaxation to acetylcholine chloride (Sigma) (10^-5^ mol/L) following a contraction evoked by phenylephrine (10^-7^ mol/L) and was defined as a relaxation ≥ 70% of the precontraction as previously described (58, 59). After extensive washout and equilibration, contraction to l-phenylephrine hydrochloride (concentration-response curve, 10−9 to 10−4 mol/L) (Sigma) and KCL (80 mM) and relaxation to acetylcholine chloride (concentration-response curve, 10−9 to 10−4 mol/L) were studied (60).

### Platelet-free plasma preparation

Beta-arrestin-2 levels were measured in the plasma of 31 patients with cirrhosis (aged 57 (47-63) years, 20 males) and 10 healthy individuals (aged 41 (34-54) years, six males). This study was approved by the Institutional Review Boards of Paris North Hospitals (Paris 7 University, France), AP-HP (N° 11-112) and Hospital Clinic (Barcelona, Spain). All patients and healthy controls included in this study gave written informed consent. The study conformed to the ethical guidelines of the 1975 Declaration of Helsinki.

Peripheral venous blood was collected from the cubital vein of fasting patients and healthy individuals, with a 21-gauge tourniquet needle, in 0.129 mol/L citrated tubes, after having discarded the first mL of blood. Tubes were then centrifuged within 2 hrs after blood draw for 15 min at 2500 g at 20°C with a light brake twice. Aliquots of platelet free plasma were then stored at −80 °C until use.

### Exosome isolation from human serum

Exosomes were immunopurified using magnetic beads (Exo-Flow, System Bioscience, Palo Alto, USA) according to the manufacturers’ protocol. Briefly, blood samples were spun at 3,000rpm for 5 min to collect the serum. The serum was then incubated directly with the magnetic beads in a 96-well plate and supernatant was decanted, while the plate was in a magnetic rack. Impurity was prevented by washing steps. Magnetic beads were removed from the exosomes with elution buffer and could be collected by decantation while the plate was in the magnetic rack.

### Determination of equilibrium AngII levels in human serum

After blood samples from hepatic vein and portal vein had been spun at 3,000rpm for 5 min, supernatant was collected, snap-frozen and stored at -80°C until measurement. Angiotensin quantification in serum was performed by RAAS equilibrium analysis in a single diagnostic liquid chromatography-mass spectrometry/mass spectroscopy (LC-MS/MS) based test by Attoquant Diagnostics (Vienna, Austria) as described previously (61, 62). Briefly, equilibrium angiotensin peptide levels were measured in conditioned and equilibrated serum samples. Subsequently, stabilized samples were spiked with stable isotope-labelled internal standards for AngII at a concentration of 200 pg/ml. Samples were subjected to LC-MS/MS analysis using a Xevo TQ-S triple quadrupole mass spectrometer (Waters, Milford, Massachusetts).

### Measurement of plasma beta-arrestin-2 levels

We determined plasma levels of beta-arrestin-2 using an ELISA kit (Human Beta-arrestin-2 ELISA Kit; # abx251362; Abbexa) according to the manufacturer’s instructions. We also determined the fraction of beta-arrestin-2 carried by microvesicles using a strategy previously described (59, 63) by measuring circulating beta-arrestin levels after filtration of the plasma through two 0.2 µm filters (Ceveron MFU 500, Technoclone, Austria). As previously described, the difference between soluble beta-arrestin levels in initial and in filtered plasma reflects the fraction of beta-arrestin carried by microvesicles.

### AT1R stimulation with AngII and SII

Persistent stimulation of the AT1R was performed with subcutaneously implanted osmotic mini-pumps (Alzet, Sulzfeld, Germany) releasing AngII or SII as described previously (5). Angiotensin II (AngII) is a member of the renin angiotensin system and an agonist of the AT1 receptor. The modified peptide [Ser(1), Ile(4), Ile(8)]-angiotensin II (SII) selectively activates the ARRB2 interaction with the AT1R. AngII and SII were released with a dose of 0.7mg/kg per day over 14 days. Animals with implanted pumps releasing an isotonic saline solvent were used as controls. For cell culture experiments, AngII or SII were added to the cell culture medium. Incubation time and dose of AngII and SII are indicated in the respective figure legends.

### Quantitative real-time polymerase chain reaction

RNA was isolated from snap-frozen samples. Reverse transcription and detection by qRT-PCR were performed following standard protocols as described previously (49, 64). The assays (Applied Biosystems, Foster City, USA) for rodent and human samples are listed in Supplemental Table 3. Primers used for experiments investigating carcinogenesis are listed in Supplemental Table 4. Results were normalized to 18sRNA, as endogenous control, expressed as 2^-ΔΔCt^ and reveal the x-fold shift of gene transcription compared to a respective control group (65).

### Transcriptome analysis of primary human HSC

Transcriptome analysis of primary human HSC was performed by OakLabs (Hennigsdorf, Germany) using Agilent Microarray XS (Agilent Technologies, Santa Clara, USA). Low Input QuickAmp Labeling Kit (Agilent Technologies, Santa Clara, USA) was used to create fluorescent complementary RNA (cRNA) followed by hybridization to microarrays using Gene Expression Hybridization Kit (Agilent Technologies, Santa Clara, USA). Fluorescence signals were detected using SureScan Microarray Scanner (Agilent Technologies, Santa Clara, USA). Transcriptome analysis of whole liver samples was performed with the Illumina NovaSeq 6000 S4 Reagent Kit (200 cycles) with the NovaSeq Xp 4-Lane Kit. The count matrix was generated using BioJupies (66), data was normalized with the *R* package edgeR (version 3.30.0) (67) and differential gene expression analysis was performed with the *R* package limma (version 3.44.1) (68). Transcriptome analysis of peripheral blood mononuclear cells was performed as described previously using Affymetrix Human Exon 1.0 ST (69). Detailed information about PBMC donors and methods can be found in the Supplemental Experimental Procedures. Gene ontology (GO) analyses of SII-dependent gene regulation was performed using Gene Ontology enRIchment anaLysis and visuaLizAtion tool (GOrilla), an online tool for discovery and visualization of enriched GO terms in ranked gene lists (http://cbl-gorilla.cs.technion.ac.il/) (70, 71). Correlation analysis of primary hHSC and whole liver samples was performed and visualized with the *R* package corrplot using the Spearman correlation method (72).

### Transcriptome analysis of human PBMC

Peripheral-blood mononuclear cells (PBMCs) were isolated from four patients with advanced alcoholic cirrhosis admitted to the Liver Unit of Beaujon Hospital, Clichy, France. Patients enrolled in the study were stable and did not have untreated or recently treated (less than one week) bacterial infection. Alcohol consumption was stopped for at least three days. In addition, four healthy subjects were investigated. The study protocol was approved by the local Institutional

Review Board and all patients signed an informed consent in accordance with the Helsinki declaration. Characteristics of patients have been previously described (69).

Affymetrix Human Exon 1.0 ST arrays were hybridized by GenoSplice technology (www.genosplice.com) according the Ambion WT protocol (Life Technologies, France) and Affymetrix (Santa Clara, CA, USA) labelling and hybridization recommendations. Affymetrix Human Exon 1.0 ST Array dataset analysis and visualization were made using EASANA® (GenoSplice Technology), which is based on the GenoSplice’s FAST DB® annotations (73, 74). An unpaired Student’s t-test was performed to compare gene intensities in the different biological replicates. Genes were considered significantly regulated when fold-change was ≥1.5 and P value ≤0.05.

### Western blotting

Snap-frozen human and murine liver samples were processed as previously described using sodium dodecyl sulfate polyacrylamide gel electrophoresis gels and nitrocellulose membranes (49). GAPDH served as endogenous control and equal protein loading was secured by Ponceau-S staining. The primary antibodies used are listed in Supplemental Table 5. Membranes were incubated with corresponding secondary peroxidase-coupled antibodies (Callbiochem, San Diego, USA). Membranes were developed with chemiluminescence (ECL, Amersham, UK) and intensities of each band were digitally detected and evaluated using Chemi-Smart (PeqLab Biotechnologies, Erlangen, Germany).

### Enzyme-linked immunosorbent assay

ELISA assays were used to measure ARRB2 (# abx251362, Abbexa, Cambridge, UK) levels in human blood samples according to the manufacturers protocol. Incubated specimens were measured at 450nm. The measured optical density was adjusted to a compiled standard curve and expressed as pg/ml. The method to determine the fraction of ARRB2 carried by microvesicles was described previously (59, 63) and can be found in the Supplemental Experimental Procedures.

### Histological staining

Visualization of collagen fibers was performed by incubation of 2-3µm thick paraffin-embedded sections with 0.1% Sirius red in saturated picric acid (Chroma, Münster, Germany) (49). Immunohistochemical staining for alpha smooth muscle actin (αSMA) was also performed on 2-3µm thick paraffin-embedded liver sections. Specimens were incubated with mouse-anti-SMA primary antibody (Actin Clone 1A4, Dako, Hamburg, Germany) and corresponding biotinylated goat-anti-mouse secondary antibody (Dako, Hamburg, Germany). Thereafter, sections were counterstained with hematoxylin (Merck, Darmstadt, Germany). Stainings were digitalized (Pannoramic MIDI) and computational analyzed (Histoquant, both 3D Histech, Budapest, Hungary) as previously described, excluding large bile ducts and vessels (5, 75). Quantification of positive staining was performed by ImageJ (version 1.51q, NIH, USA).

### Tissue micro array

Tissue micro arrays were prepared using paired HCC and non-neoplastic liver tissues from HCC patients undergoing tumor resection as described previously (23). Specimens were incubated with rabbit-anti-ARRB2 primary antibody (Cell Signaling Technology, USA) or rabbit-anti-RARRES1 primary antibody (Abcam, Cambridge, UK) and corresponding HRP labelled polymer (goat-anti-rabbit secondary antibody). Thereafter, sections were counterstained with hematoxylin (Merck, Darmstadt, Germany). All patients signed an informed consent in accordance with the declaration of Helsinki.

### *In silico* validation

The Oncomine cancer microarray database (https://www.oncomine.org/) was used to analyze *ARRB2* and *RARRES1* expression in diverse datasets (15, 76) as described elsewhere (23, 77). The impact of *ARRB2* and *RARRES1* expression on survival in HCC was analyzed in The Cancer Genome Atlas (TCGA) dataset using the online biomarker validation tool SurvExpress as described previously (23, 78). Expression of *ARRB2* and *RARRES1* in murine HCC and non-HCC controls was analyzed using the GEO datasets (GEO profiles GDS3087/1426239_s_at, GDS3087/1451987_at, GDS3087 / 1438055_at).

### Statistics

Statistical analysis was performed with Prism (GraphPad, La Jolla, USA). Data are presented as mean ± standard error of the mean (SEM). Real-time PCR and ELISA were performed in duplicates. Cell culture experiments were replicated three times from at least three different isolations. The *n* for each experiment is given, when not represented graphically in the figure. Differences between the groups were analyzed using the nonparametric Mann-Whitney-U test or using the Wilcoxon matched pairs test when values represent paired observations. For transcriptome analysis the Welch’s t-test was used and *p*-values were adjusted for multiple testing using the Benjamini–Hochberg FDR procedure (79). A *p*-value < 0.05 was considered statistically significant. Analysis of tissue micro arrays was performed applying the Fisher’s exact test and Spearman correlation analysis. *In silico* survival analysis were performed computationally using Log-rank testing and Hazard ratio estimates.

### Study approval

The use of human liver samples was approved by the Human Ethics Committee of the University of Bonn (file reference: 202/01 and 029/13). Sampling of human livers for primary cell isolation was approved by the Ethics Committee of the University College London (NRES Rec Reference NC2015.020). All patients signed an informed consent in accordance with the Helsinki declaration before being enrolled in the study. All animal experiments were performed in accordance with the German animal protection law and the guidelines of the animal care unit at the University of Bonn (Haus für experimentelle Therapie, University Hospital Bonn, Germany) and the Goethe University Frankfurt (Zentrale Forschungseinrichtung, Goethe University Frankfurt, Germany) and approved by the relevant state agencies (file reference: LANUV 84-02.04.2015.A491, 84-02.04.2013.A129 and FK/1119).

### Funding

German Research Foundation grant SFB TRR57 (JT)

German Research Foundation grant CRC1382 (JT)

Gisela Stadelmann-Foundation of the University Hospital Frankfurt (RS)

August Scheidel-Foundation of the University Hospital Frankfurt (RS)

Martha Schmelz estate managed by the University Hospital Frankfurt (RS)

Eurostars grant 12350 (JT)

European Union’s Horizon 2020 research and innovation programme MICROB-PREDICT, grant 825694 (JT)

European Union’s Horizon 2020 research and innovation programme GALAXY, grant 668031 (JT)

European Union’s Horizon 2020 research and innovation programme LIVERHOPE, grant 731875 (JT)

The manuscript reflects only the author’s view, and the European Commission is not responsible for any use that may be made of the information it contains. The funders had no influence on study design, data collection and analysis, decision to publish or preparation of the manuscript.

## Author contributions

Conceptualization and methodology: R.S. and J.T.

Investigation: R.S., J.H., P.D., S.K., F.E.U., C.O., O.T., S.T., Ch.H., N.K., R.P., J.P., P.E.R., G.K., J.G.S., M.P., I.L.P., T.W., C.T., Z.A., C.M., M.F., R.M., Cl.H., K.R., W.K., A.M.D. and J.T.

Resources: P.D., L.D.G., G.K., K.R, W.K., A.M.D, M.P., T.W., R.M., Cl.H., P.E.R. and J.T.

Writing – original draft: R.S. and J.T.

Writing – review & editing: R.S., J.H., P.D., S.K., F.E.U., C.O., O.T., S.T., Ch.H., N.K., R.P., L.D.G., J.P., P.E.R., G.K., J.G.S., M.P., I.L.P., T.W., C.T., Z.A., C.M., M.F., R.M., Cl.H., K.R., W.K., A.M.D. and J.T.

Visualization: R.S., P.D., S.K., P.E.R., J.G.S.

Funding acquisition: J.T., R.S. Supervision: J.T.

## Acknowledgments

The authors thank Gudrun Hack, Silke Bellinghausen, Evelyn Süss, Dikra Zouiten, Christian Schmithals and Maike von Harten for excellent technical assistance. The authors thank Nikos Werner and Robert Lefkowitz for the provision of *Agtr1*^-/-^ and *Arrb2*^-/-^ mice.

## Supplementary Materials

### Supplementary figures

Fig. S1.

Fig. S2.

Fig. S3.

Fig. S4.

Fig. S5.

### Supplementary tables

Table S1. Gene ontology enrichment analysis of upregulated genes.

Table S2. Gene ontology enrichment analysis of downregulated genes.

Table S3. Taqman assays.

Table S4. Primer list.

Table S5. Primary antibodies for Western blot, immunohistochemistry and tissue micro array.

## References

1. Wynn TA, Ramalingam TR. Mechanisms of fibrosis: therapeutic translation for fibrotic disease. Nat Med 2012;18(7):1028–1040.

2. Tsuchida T, Friedman SL. Mechanisms of hepatic stellate cell activation. Nat Rev Gastroenterol Hepatol 2017;14(7):397–411.

3. Teufel A et al. Identification of RARRES1 as a core regulator in liver fibrosis. J. Mol. Med. 2012;90(12):1439–1447.

4. Mederacke I et al. Fate tracing reveals hepatic stellate cells as dominant contributors to liver fibrosis independent of its aetiology [Internet]. Nat Commun 2013;4. doi:10.1038/ncomms3823

5. Granzow M et al. Angiotensin-II type 1 receptor-mediated Janus kinase 2 activation induces liver fibrosis. Hepatology 2014;60(1):334–348.

6. Klein S et al. Janus-kinase-2 relates directly to portal hypertension and to complications in rodent and human cirrhosis. Gut 2017;66(1):145–155.

7. Gurevich VV, Gurevich EV. The structural basis of arrestin-mediated regulation of G-protein-coupled receptors. Pharmacol Ther 2006;110(3):465–502.

8. Lovgren AK et al. β-arrestin deficiency protects against pulmonary fibrosis in mice and prevents fibroblast invasion of extracellular matrix. Sci Transl Med 2011;3(74):74ra23.

9. Gurevich VV, Gurevich EV. GPCR Signaling Regulation: The Role of GRKs and Arrestins. Front Pharmacol 2019;10:125.

10. Kim J et al. Beta-arrestins regulate atherosclerosis and neointimal hyperplasia by controlling smooth muscle cell proliferation and migration. Circ. Res. 2008;103(1):70–79.

11. Yang Y et al. β-Arrestin1 enhances hepatocellular carcinogenesis through inflammation-mediated Akt signalling. Nat Commun 2015;6:7369.

12. Rajagopal K et al. Beta-arrestin2-mediated inotropic effects of the angiotensin II type 1A receptor in isolated cardiac myocytes. Proc. Natl. Acad. Sci. U.S.A. 2006;103(44):16284–16289.

13. Schierwagen R et al. β-Arrestin2 is increased in liver fibrosis in humans and rodents. PNAS 2020;117(44):27082–27084.

14. Ranjan R, Gupta P, Shukla AK. GPCR Signaling: β-arrestins Kiss and Remember. Current Biology 2016;26(7):R285–R288.

15. Mas VR et al. Genes involved in viral carcinogenesis and tumor initiation in Hepatitis C Virus-induced Hepatocellular Carcinoma. Mol Med 2009;15(3–4):85–94.

16. Piu F, Gauthier NK, Wang F. Beta-arrestin 2 modulates the activity of nuclear receptor RAR beta2 through activation of ERK2 kinase. Oncogene 2006;25(2):218–229.

17. Nagpal S et al. Tazarotene-induced gene 1 (TIG1), a novel retinoic acid receptor-responsive gene in skin. J Invest Dermatol 1996;106(2):269–274.

18. De Minicis S et al. Gene expression profiles during hepatic stellate cell activation in culture and in vivo. Gastroenterology 2007;132(5):1937–1946.

19. Trebicka J. Emergency TIPS in a Child-Pugh B patient: When does the window of opportunity open and close?. J. Hepatol. 2017;66(2):442–450.

20. Pironti G et al. Circulating Exosomes Induced by Cardiac Pressure Overload Contain Functional Angiotensin II Type 1 Receptors. Circulation 2015;131(24):2120–2130.

21. Chen L, Chen R, Velazquez VM, Brigstock DR. Fibrogenic Signaling Is Suppressed in Hepatic Stellate Cells through Targeting of Connective Tissue Growth Factor (CCN2) by Cellular or Exosomal MicroRNA-199a-5p. Am. J. Pathol. 2016;186(11):2921–2933.

22. Charrier A et al. Exosomes mediate intercellular transfer of pro-fibrogenic connective tissue growth factor (CCN2) between hepatic stellate cells, the principal fibrotic cells in the liver. Surgery 2014;156(3):548–555.

23. Dietrich P et al. Wild type Kirsten rat sarcoma is a novel microRNA-622-regulated therapeutic target for hepatocellular carcinoma and contributes to sorafenib resistance. Gut 2018;67(7):1328–1341.

24. Bataller R et al. Angiotensin II induces contraction and proliferation of human hepatic stellate cells. Gastroenterology 2000;118(6):1149–1156.

25. Bataller R et al. NADPH oxidase signal transduces angiotensin II in hepatic stellate cells and is critical in hepatic fibrosis. Journal of Clinical Investigation 2003;112(9):1383–1394.

26. Kendall RT et al. The beta-arrestin pathway-selective type 1A angiotensin receptor (AT1A) agonist [Sar1,Ile4,Ile8]angiotensin II regulates a robust G protein-independent signaling network. J. Biol. Chem. 2011;286(22):19880–19891.

27. Sun W-Y et al. Depletion of β-arrestin2 in hepatic stellate cells reduces cell proliferation via ERK pathway. J. Cell. Biochem. 2013;114(5):1153–1162.

28. Li J et al. β-Arrestins regulate human cardiac fibroblast transformation and collagen synthesis in adverse ventricular remodeling. J. Mol. Cell. Cardiol. 2014;76:73–83.

29. Klein S et al. HSC-specific inhibition of Rho-kinase reduces portal pressure in cirrhotic rats without major systemic effects. Journal of Hepatology 2012;57(6):1220–1227.

30. Klein S et al. Rho-kinase inhibitor coupled to peptide-modified albumin carrier reduces portal pressure and increases renal perfusion in cirrhotic rats. Sci Rep 2019;9(1):2256.

31. Kotula JW et al. Targeted disruption of β-arrestin 2-mediated signaling pathways by aptamer chimeras leads to inhibition of leukemic cell growth. PLoS ONE 2014;9(4):e93441.

32. Yin D et al. β-Arrestin 2 Promotes Hepatocyte Apoptosis by Inhibiting Akt Protein. J. Biol. Chem. 2016;291(2):605–612.

33. Gonzalez-Begne M et al. Proteomic Analysis of Human Parotid Gland Exosomes by Multidimensional Protein Identification Technology (MudPIT). Journal of proteome research 2009;8(3):1304.

34. Hennenberg M et al. Vascular dysfunction in human and rat cirrhosis: Role of receptor-desensitizing and calcium-sensitizing proteins. Hepatology 2007;45(2):495–506.

35. Liu S, Luttrell LM, Premont RT, Rockey DC. β-Arrestin2 is a critical component of the GPCR–eNOS signalosome. PNAS 2020;117(21):11483–11492.

36. Chen X-H, Wu W-G, Ding J. Aberrant TIG1 methylation associated with its decreased expression and clinicopathological significance in hepatocellular carcinoma. Tumour Biol. 2014;35(2):967–971.

37. Kloth M et al. The SNP rs6441224 influences transcriptional activity and prognostically relevant hypermethylation of RARRES1 in prostate cancer. Int. J. Cancer 2012;131(6):E897–904.

38. Trebicka J et al. Role of beta3-adrenoceptors for intrahepatic resistance and portal hypertension in liver cirrhosis. Hepatology 2009;50(6):1924–1935.

39. Nielsen MJ et al. Circulating Elastin Fragments Are Not Affected by Hepatic, Renal and Hemodynamic Changes, But Reflect Survival in Cirrhosis with TIPS. Dig. Dis. Sci. 2015;60(11):3456–3464.

40. Trebicka J et al. Endotoxin and tumor necrosis factor-receptor levels in portal and hepatic vein of patients with alcoholic liver cirrhosis receiving elective transjugular intrahepatic portosystemic shunt:. European Journal of Gastroenterology & Hepatology 2011;23(12):1218– 1225.

41. Burson JM, Aguilera G, Gross KW, Sigmund CD. Differential expression of angiotensin receptor 1A and 1B in mouse. Am. J. Physiol. 1994;267(2 Pt 1):E260–267.

42. Klein S, Schierwagen R, Uschner FE, Trebicka J. Mouse and Rat Models of Induction of Hepatic Fibrosis and Assessment of Portal Hypertension. Methods Mol. Biol. 2017;1627:91–116.

43. Domenicali M et al. A novel model of CCl_4_-induced cirrhosis with ascites in the mouse. J. Hepatol. 2009;51(6):991–999.

44. Uehara T, Pogribny IP, Rusyn I. The DEN and CCl_4_ -Induced Mouse Model of Fibrosis and Inflammation-Associated Hepatocellular Carcinoma. Curr Protoc Pharmacol 2014;66:14.30.1– 10.

45. Höchst B et al. Differential Induction of Ly6G and Ly6C Positive Myeloid Derived Suppressor Cells in Chronic Kidney and Liver Inflammation and Fibrosis [Internet]. PLoS One 2015;10(3). doi:10.1371/journal.pone.0119662

46. Tamura M, Aizawa R, Hori M, Ozaki H. Progressive renal dysfunction and macrophage infiltration in interstitial fibrosis in an adenine-induced tubulointerstitial nephritis mouse model. Histochem Cell Biol 2009;131(4):483–490.

47. Haupenthal J et al. Reduced Efficacy of the Plk1 Inhibitor BI 2536 on the Progression of Hepatocellular Carcinoma due to Low Intratumoral Drug Levels. Neoplasia 2012;14(5):410– 419.

48. Schmithals C et al. Improving Drug Penetrability with iRGD Leverages the Therapeutic Response to Sorafenib and Doxorubicin in Hepatocellular Carcinoma. Cancer Res 2015;75(15):3147–3154.

49. Trebicka J et al. Atorvastatin lowers portal pressure in cirrhotic rats by inhibition of RhoA/Rho-kinase and activation of endothelial nitric oxide synthase. Hepatology 2007;46(1):242–253.

50. Rombouts K, Carloni V. Determination and Characterization of Tetraspanin-Associated Phosphoinositide-4 Kinases in Primary and Neoplastic Liver Cells. Methods Mol. Biol. 2016;1376:203–212.

51. Ortega-Ribera M et al. Resemblance of the human liver sinusoid in a fluidic device with biomedical and pharmaceutical applications. Biotechnol Bioeng 2018;115(10):2585–2594.

52. Wojtalla A et al. The endocannabinoid N-arachidonoyl dopamine (NADA) selectively induces oxidative stress-mediated cell death in hepatic stellate cells but not in hepatocytes. Am. J. Physiol. Gastrointest. Liver Physiol. 2012;302(8):G873–887.

53. Schiwon M et al. Crosstalk between Sentinel and Helper Macrophages Permits Neutrophil Migration into Infected Uroepithelium. Cell 2014;156(3):456–468.

54. Kastenmüller W, Torabi-Parizi P, Subramanian N, Lämmermann T, Germain RN. A spatially-organized multicellular innate immune response in lymph nodes limits systemic pathogen spread. Cell 2012;150(6):1235–1248.

55. Kastenmüller W et al. Peripheral pre-positioning and local CXCL9 chemokine-mediated guidance orchestrate rapid memory CD8+ T cell responses in the lymph node. Immunity 2013;38(3):502–513.

56. Lee SML, Schelcher C, Demmel M, Hauner M, Thasler WE. Isolation of human hepatocytes by a two-step collagenase perfusion procedure. J Vis Exp [published online ahead of print: September 3, 2013];(79). doi:10.3791/50615

57. Bauer R et al. Downregulation of P-cadherin expression in hepatocellular carcinoma induces tumorigenicity. Int J Clin Exp Pathol 2014;7(9):6125–6132.

58. Poisson J et al. Erythrocyte-derived microvesicles induce arterial spasms in JAK2V617F myeloproliferative neoplasm. J. Clin. Invest. [published online ahead of print: February 11, 2020]; doi:10.1172/JCI124566

59. Rautou P et al. Abnormal Plasma Microparticles Impair Vasoconstrictor Responses in Patients With Cirrhosis. Gastroenterology 2012;143(1):166–176.e6.

60. Heymes C et al. Cyclo-oxygenase-1 and -2 contribution to endothelial dysfunction in ageing. Br. J. Pharmacol. 2000;131(4):804–810.

61. Basu R et al. Roles of Angiotensin Peptides and Recombinant Human ACE2 in Heart Failure. J. Am. Coll. Cardiol. 2017;69(7):805–819.

62. Pavo N et al. Low- and High-renin Heart Failure Phenotypes with Clinical Implications. Clin. Chem. 2018;64(3):597–608.

63. Bissonnette J et al. A prospective study of the utility of plasma biomarkers to diagnose alcoholic hepatitis. Hepatology 2017;66(2):555–563.

64. Schierwagen R et al. Seven weeks of Western diet in apolipoprotein-E-deficient mice induce metabolic syndrome and non-alcoholic steatohepatitis with liver fibrosis. Sci Rep 2015;5:12931.

65. Livak KJ, Schmittgen TD. Analysis of relative gene expression data using real-time quantitative PCR and the 2(-Delta Delta C(T)) Method. Methods 2001;25(4):402–408.

66. Torre D, Lachmann A, Ma’ayan A. BioJupies: Automated Generation of Interactive Notebooks for RNA-Seq Data Analysis in the Cloud. Cell Syst 2018;7(5):556–561.e3.

67. Robinson MD, McCarthy DJ, Smyth GK. edgeR: a Bioconductor package for differential expression analysis of digital gene expression data. Bioinformatics 2010;26(1):139–140.

68. Ritchie ME et al. limma powers differential expression analyses for RNA-sequencing and microarray studies. Nucleic Acids Res 2015;43(7):e47.

69. Gandoura S et al. Gene- and exon-expression profiling reveals an extensive LPS-induced response in immune cells in patients with cirrhosis. J. Hepatol. 2013;58(5):936–948.

70. Eden E, Lipson D, Yogev S, Yakhini Z. Discovering motifs in ranked lists of DNA sequences. PLoS Comput. Biol. 2007;3(3):e39.

71. Eden E, Navon R, Steinfeld I, Lipson D, Yakhini Z. GOrilla: a tool for discovery and visualization of enriched GO terms in ranked gene lists. BMC Bioinformatics 2009;10:48.

72. Wei T, Simko V. corrplot: Visualization of a correlation matrix. R package version 0.73 2013;230(231):11.

73. de la Grange P, Dutertre M, Martin N, Auboeuf D. FAST DB: a website resource for the study of the expression regulation of human gene products. Nucleic Acids Res. 2005;33(13):4276–4284.

74. de la Grange P, Dutertre M, Correa M, Auboeuf D. A new advance in alternative splicing databases: from catalogue to detailed analysis of regulation of expression and function of human alternative splicing variants. BMC Bioinformatics 2007;8:180.

75. Huss S et al. Development and evaluation of an open source Delphi-based software for morphometric quantification of liver fibrosis. Fibrogenesis Tissue Repair 2010;3:10.

76. Su AI et al. Molecular classification of human carcinomas by use of gene expression signatures. Cancer Res. 2001;61(20):7388–7393.

77. Rhodes DR et al. ONCOMINE: A Cancer Microarray Database and Integrated Data-Mining Platform. Neoplasia 2004;6(1):1–6.

78. Aguirre-Gamboa R et al. SurvExpress: an online biomarker validation tool and database for cancer gene expression data using survival analysis. PLoS ONE 2013;8(9):e74250.

79. Benjamini Y, Hochberg Y. Controlling the False Discovery Rate: A Practical and Powerful Approach to Multiple Testing. Journal of the Royal Statistical Society. Series B (Methodological) 1995;57(1):289–300.

