## Supplementary Material for "Intercellular crosstalk regulating ARRB2/RARRES1 is involved in transition from fibrosis to cancer"

### Supplemental data

#### Supplemental Table 1: Gene ontology enrichment analysis of upregulated genes

| Gene Ontology term | gene count | fold enrichment | *p* value |
|---|---|---|---|
| BP: interferon-gamma-mediated signaling pathway | 16 | 7.89 | 9.18E-10 |
| BP: regulation of interleukin-1 beta production | 11 | 6.04 | 7.74E-06 |
| CC: clathrin-coated endocytic vesicle membrane | 12 | 5.36 | 6.00E-06 |
| BP: positive regulation of interleukin-6 production | 17 | 4.06 | 2.00E-06 |
| BP: regulation of cartilage development | 16 | 3.93 | 4.74E-06 |
| BP: regulation of receptor activity | 24 | 3.09 | 6.55E-06 |
| MF: growth factor binding | 26 | 3.03 | 1.14E-06 |
| BP: response to external biotic stimulus | 65 | 2.61 | 6.74E-12 |
| BP: cytokine-mediated signaling pathway | 64 | 2.49 | 4.88E-11 |
| BP: positive regulation of cell activation | 37 | 2.37 | 3.69E-06 |
| BP: positive regulation of cell migration | 62 | 2.25 | 3.30E-09 |
| CC: extracellular space | 68 | 2.19 | 6.63E-09 |
| CC: cell surface | 70 | 2.03 | 3.01E-08 |
| BP: positive regulation of cell differentiation | 91 | 1.92 | 4.98E-09 |
| BP: regulation of MAPK cascade | 59 | 1.90 | 9.89E-06 |
| BP: positive regulation of cell proliferation | 71 | 1.88 | 1.12E-06 |
| BP: cell surface receptor signaling pathway | 162 | 1.70 | 2.86E-11 |
| BP: regulation of programmed cell death | 120 | 1.57 | 2.72E-06 |
| BP: regulation of signal transduction | 111 | 1.56 | 8.12E-06 |
| BP: positive regulation of cell communication | 142 | 1.49 | 4.09E-06 |

#### Supplemental Table 2: Gene ontology enrichment analysis of downregulated genes

| Gene Ontology term | gene count | fold enrichment | *p* value |
|---|---|---|---|
| BP: extracellular matrix disassembly | 6 | 24.16 | 1.68E-06 |
| MF: cytokine receptor binding | 9 | 10.37 | 2.77E-06 |
| MF: receptor regulator activity | 14 | 9.89 | 5.57E-10 |
| MF: receptor ligand activity | 13 | 9.67 | 3.79E-09 |
| BP: regulation of receptor activity | 13 | 8.21 | 2.99E-08 |
| MF: chemokine receptor binding | 11 | 8.07 | 5.89E-07 |
| CC: extracellular space | 25 | 3.99 | 1.15E-08 |
| BP: cellular response to lipid | 25 | 3.09 | 3.14E-06 |
| BP: regulation of cell proliferation | 46 | 2.25 | 1.75E-06 |
| BP: regulation of response to stimulus | 94 | 1.72 | 2.40E-07 |

#### Supplemental Table 3: Taqman assays

| Gene | Assay ID | Species |
|---|---|---|
| *Acta2* | Mm00725412_s1 | Mus musculus |
| *Agtr1* | Mm01166161_m1 | Mus musculus |
| *ARRB2* | Hs00188826_m1 | Human |

| | | |
|---|---|---|
| *Arrb2* | Mm00520665-m1 | Mus musculus |
| *Arrb2* | Rn00563775_m1 | Rattus norvegicus |
| *Col1a1* | Mm00801666_g1 | Mus musculus |
| *RARRES1* | Hs00894859_m1 | Human |
| *Rarres1* | Mm01220691_m1 | Mus musculus |

**Supplemental Table 4: Primer list**

| **Gene: *Arrb2*** | **Species: Mus musculus** |
|---|---|
| forward | CGTCTGTCCACCCGAGATAC |
| reverse | CGAGTTGGTGTGAGGCCAAA |
| **Gene: *ARRB2*** | **Species: human** |
| forward | CTGTAGATGGCGTGGTGCTT |
| reverse | TGGTAGGTGGCGATGAACAG |
| **Gene: *Rarres1*** | **Species: Mus musculus** |
| forward | CCTTCCTCGGCAGCTCATAC |
| reverse | ACCAAGTGAATACGGCAGGG |
| **Gene: *RARRES1*** | **Species: human** |
| forward | CGCTACAACCCAGAGTCTTTACT |
| reverse | AGCAGGTAATCCTCTTGTTGTCTT |

**Supplemental Table 5: Primary antibodies for Western blot, immunohistochemistry and tissue micro array**

| **Name** | **Order number** | **Company** |
|---|---|---|
| ARRB2 | ab54790 | Abcam plc, Cambridge, UK |
| ARRB2 | 3857 | Cell Signaling Technology Inc., MA, USA |
| Alpha-SMA | M0851 | Dako, Hamburg, Germany |
| ERK1/2 | sc-93 | Santa Cruz Biotechnology, Santa Cruz, CA, USA |
| GAPDH | sc-25778 | Santa Cruz Biotechnology, Santa Cruz, CA, USA |
| pERK1/2 | 4370 | Cell Signaling Technology Inc., MA, USA |
| RARRES1 | ab198908 | Abcam plc, Cambridge, UK |

### Supplemental figures

#### Supp. Figure 1

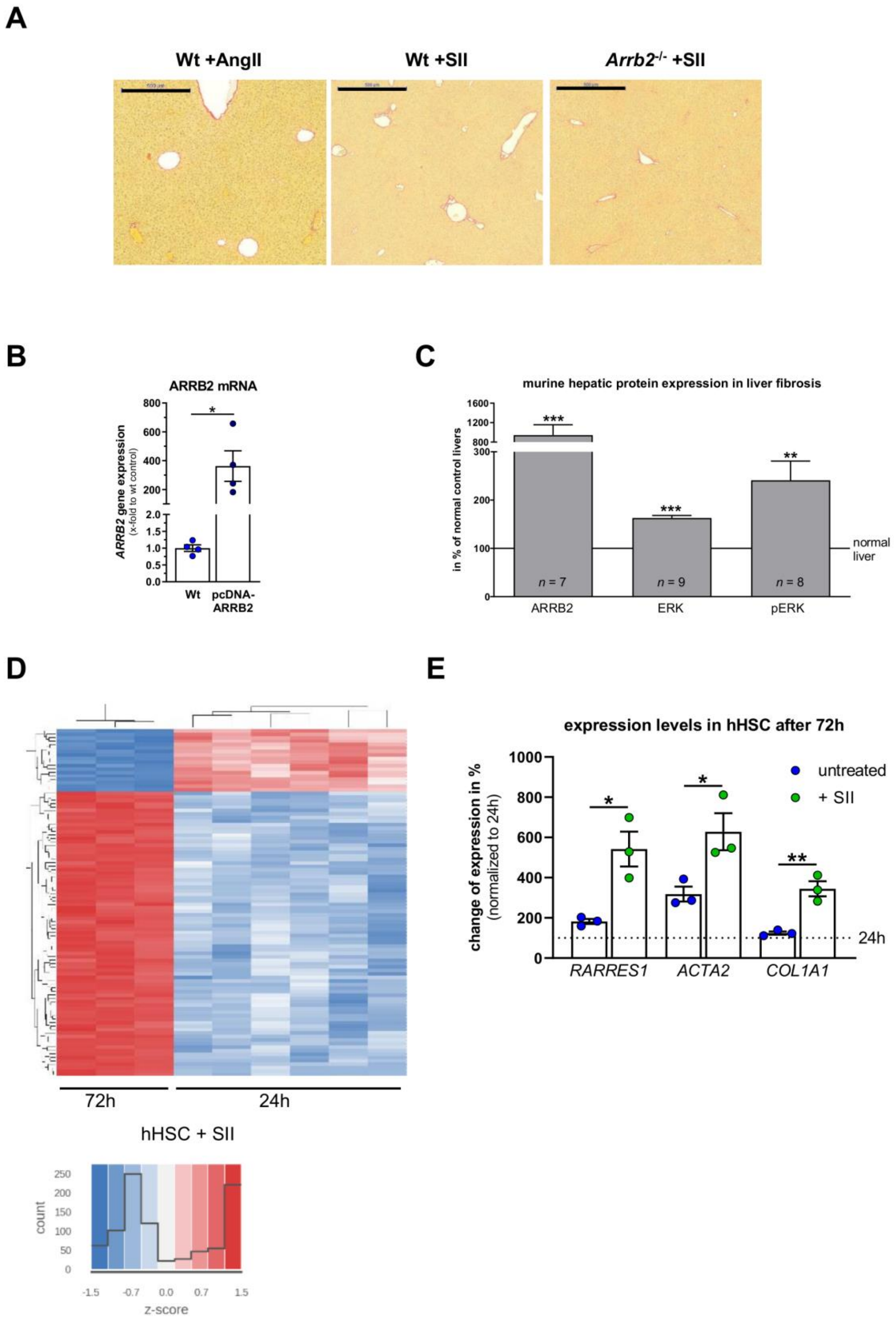

**Supplemental Figure 1.** (A) Representative Sirius red stainings of Wt and beta-arrestin-2 deficient (*Arrb2*$^{-/-}$) mice after angiotensin II receptor type I subtype A (AT1R) stimulation either G-protein-dependent by angiotensin II (AngII) or G-protein-independent via beta-arrestin-2 (ARRB2) by the modified peptide [Ser(1), Ile(4), Ile(8)]-angiotensin II (SII) for 14 days using osmotic mini-pumps. Scale bars are 500µm. (B) *ARRB2* expression in LX2 cells transfected with ARRB2 plasmids or control plasmids. (C) Quantification of ARRB2 and downstream protein expression from Western blot in murine liver tissue after fibrosis induction using carbon tetrachloride ($CCl_4$) intoxication three times a week for four weeks or by bile duct ligation (BDL) compared to untreated control animals. (D) Transcriptome analyses were performed by RNA microarray in primary human HSC (hHSC) incubated with SII for 24 hrs (n=6) or 72 hrs (n=3) to elucidate time-dependent effects of ARRB2 stimulation at the stage of early and full HSC activation. Blue: downregulated genes, red: upregulated genes. (E) Change of gene expression in untreated or SII treated hHSC harvested after 72h in relation to expression of cells harvested after 24h (dotted line). (Graphs show bar graphs with mean ± SEM and *n* value given for each group. * = $p < 0.05$, ** = $p < 0.01$, *** = $p < 0.001$.

Supp. Figure 2

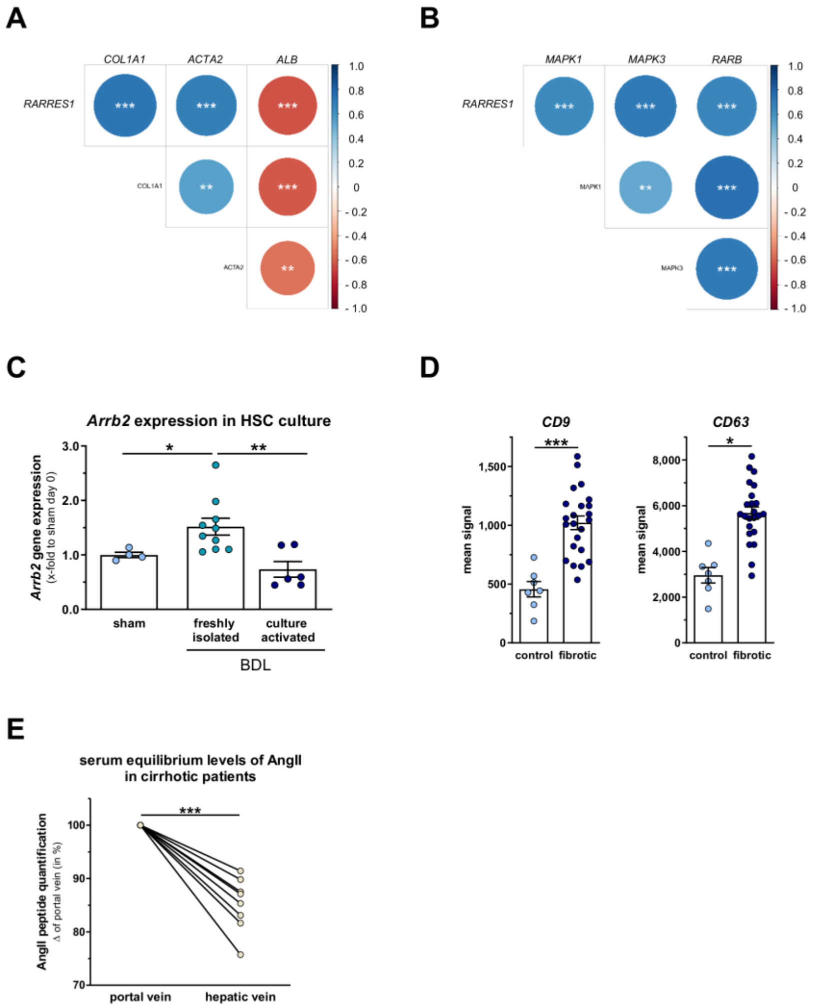


**Supplemental Figure 2.** (A) Correlation of *RARRES1* with markers of downstream signaling via extracellular-signal regulated kinase 1/2 (*ERK1/2*) (*MAPK1*, *MAPK3*), hepatic stellate cell (HSC) activity markers (*COL1A1*, *ACTA2*) and ALB for hepatocyte function in transcriptome data from whole liver tissue. (B) Correlation of RARRES1 with markers of downstream signaling via extracellular-signal regulated kinase 1/2 (MAPK1, MAPK3) and retinoic acid receptor beta (RARB). (C) *ARRB2* expression in freshly isolated or culture activated hepatic stellate cells (HSC) compared to HSC from sham operated rats. (D) Expression of exosome markers (*CD9*, *CD63*) in transcriptome data from whole liver tissue of fibrotic and non-fibrotic individuals. (E) Serum equilibrium levels of AngII in matched portal and hepatic vein samples from cirrhotic patients. Graphs show single measurements as dots and mean ± SEM. * = $p < 0.05$, ** = $p < 0.01$, *** = $p < 0.001$.

#### Supp. Figure 3

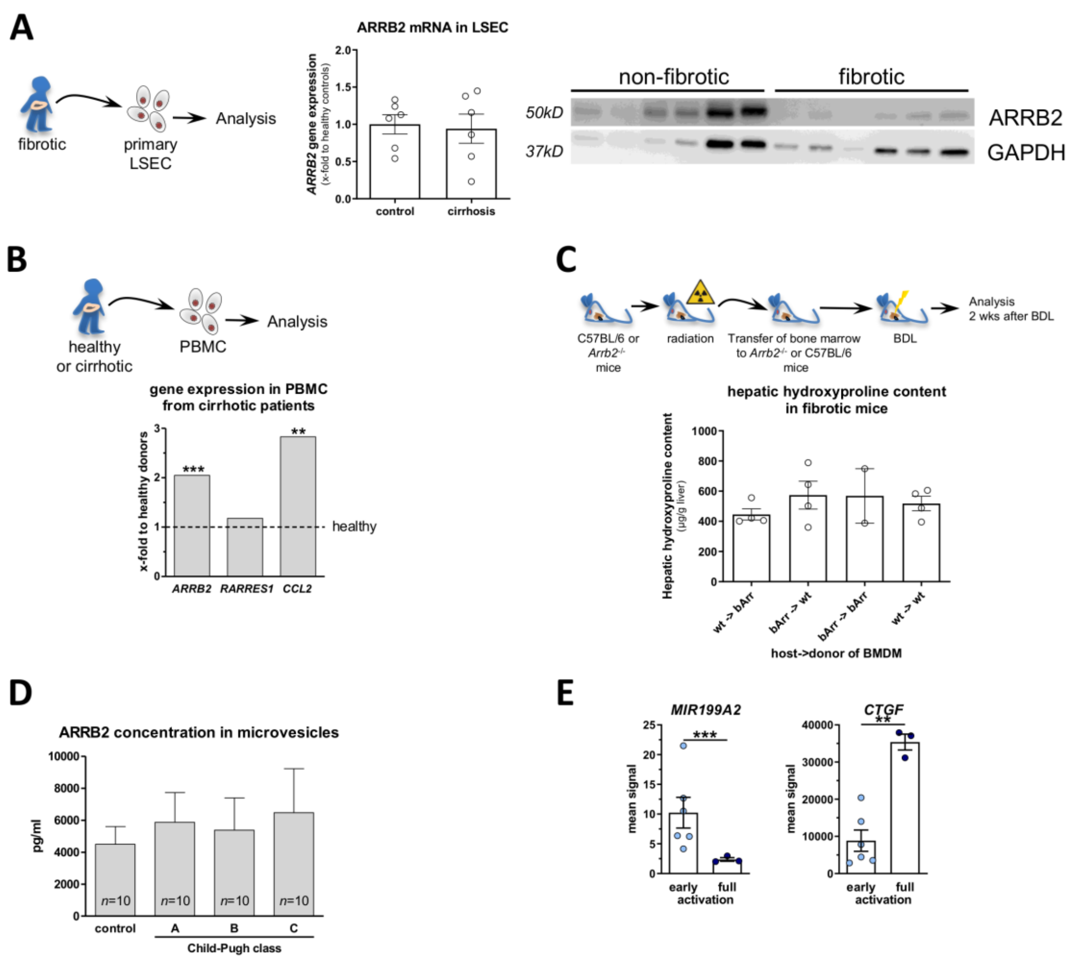


**Supplemental Figure 3.** (A) Quantification of ARRB2 gene and protein expression in primary human liver sinusoidal endothelial cells (LSEC) isolated from non-cirrhotic control and cirrhotic liver samples. (B) Results of *ARRB2*, *RARRES1* and *CCL2* mRNA expressions obtained using microarrays in peripheral-blood mononuclear cells (PBMCs) from patients with cirrhosis and healthy subjects. (C) Measurement of hepatic hydroxyproline content in Wt or *Arrb2*$^{-/-}$ mice after generation of bone marrow chimera and macrophage transfer. (D) Quantification of circulating ARRB2-loaded microvesicles in healthy control individuals and cirrhotic patients stratified by Child-Pugh classes A-C. (E) Expression of MIR199A2 and CTGF in human primary HSC harvested after 24 h (early activation) or 72 h (full activation) of incubation with SII. Graphs show bar graphs with mean ± SEM and *n* value given for each group. ** = $p < 0.01$, *** = $p < 0.001$.

Supp. Figure 4

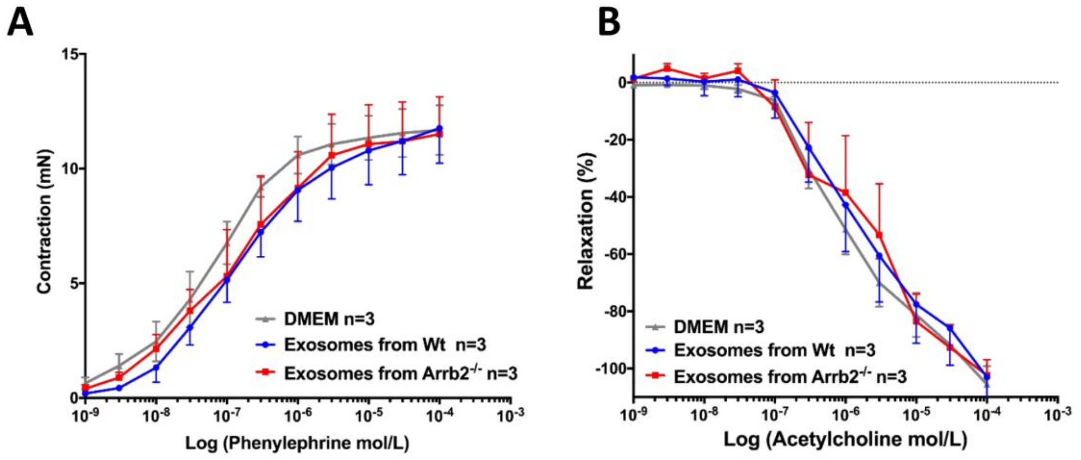


**Supplemental Figure 4.** Effect of exosomes isolated from Wt or *Arrb2*$^{-/-}$ mice on (A) aortic ring contraction and (B) aortic ring relaxation. Graphs show bar graphs with mean ± SEM.

#### Supp. Figure 5

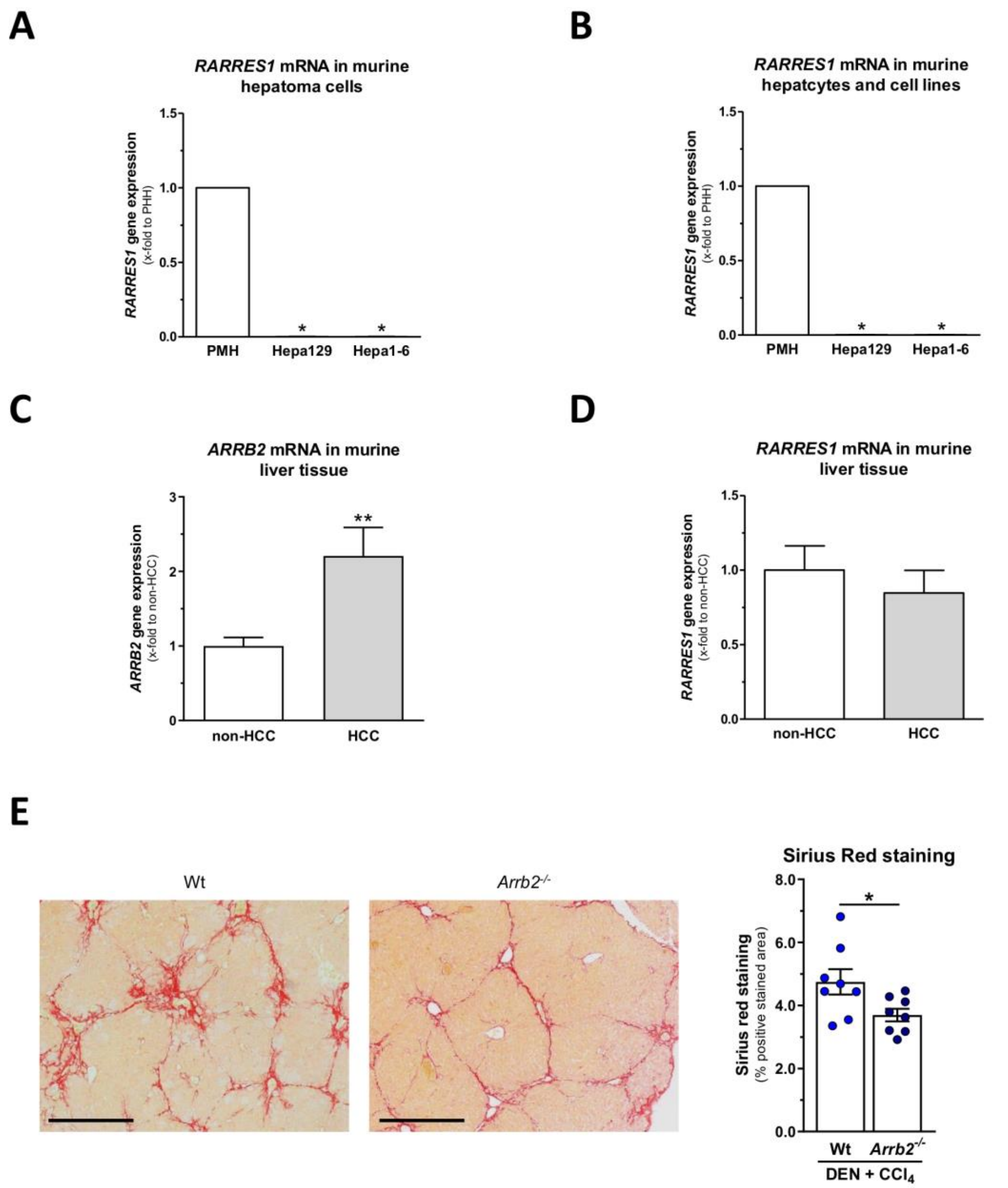


**Supplemental Figure 5.** (A) Quantification of *RARRES1* expression in primary human hepatocytes (PHH) and hepatoma cell lines HepG2 and PLC. (B) Quantification of *Rarres1* expression in primary mouse hepatocytes (PMH) and hepatoma cell lines Hepa129 and Hepa1-6. (C) *Arrb2* expression in murine non-HCC and HCC liver tissue. (D) *Rarres1* expression in murine non-HCC and HCC liver tissue. (E) Representative sections and quantification of liver fibrosis by Sirius red staining of livers from Wt and *Arrb2*$^{-/-}$ mice after single DEN injection and repetitive CCl4 intoxication. Graphs show bar graphs with mean ± SEM. $^{\#}$ = $p$ 0.1 - 0.05, * = $p < 0.05$, ** = $p < 0.01$.
